# A short course of antibiotics selects for persistent resistance in the human gut

**DOI:** 10.1101/2023.09.04.556257

**Authors:** Eitan Yaffe, Les Dethlefsen, Arati V. Patankar, Chen Gui, Susan Holmes, David A. Relman

## Abstract

Understanding the relationship between antibiotic use and the evolution of antimicrobial resistance is vital for effective antibiotic stewardship, yet animal models and *in vitro* experiments poorly replicate real-world conditions. To elucidate how resistance evolves *in vivo*, we exposed 60 human subjects to ciprofloxacin and used longitudinal stool samples and a new computational method to assemble genomes for 5665 populations of commensal bacterial species within subjects. Analysis of 2.27M polymorphic sequence variants revealed 513 populations that underwent selective sweeps. We found convergent evolution focused on DNA gyrase and evidence of dispersed selective pressure at other genomic loci. Nearly 10% of susceptible bacterial populations evolved towards resistance through sweeps that involved mutations in a specific amino acid in gyrase. Evolution towards resistance was predicted by population abundances before and during the exposure. 89% of gyrase sweeps and the majority of all sweeps persisted more than 10 weeks. This work quantifies the direct relationship between antibiotic usage and the evolution of resistance within the gut communities of individual human hosts.

## INTRODUCTION

Antimicrobial resistance (AMR) is a growing global health threat, projected to cause millions of deaths worldwide by 2050^1,2^. AMR is driven by the overuse and misuse of antibiotics^3^, and is associated with geographic factors^4^, seasonal patterns^5^, and prolonged antibiotic usage^6^. AMR is not limited to heavily prescribed patients and is instead linked to low-intensity outpatient antibiotic usage^7^. Prescription data suggest that AMR primarily evolves due to bystander selective pressure, where colonizing pathogens evolve resistance subclinically in patients taking antibiotics for unrelated problems^8^. However, the evolution of AMR *in vivo* is rarely documented; instead, resistant pathogens usually emerge in infections after evolving elsewhere, as seen for example with resistant *Escherichia coli* associated with urinary tract infections^9^ and surgical site infections^10^. In contrast to the sparsity of data from treated human subjects, the evolution of AMR has been extensively studied *in vitro*^11–17^ and using animal models^18,19^. Yet, these laboratory experiments struggle to reproduce clinical conditions faithfully, most of which are poorly understood, including local drug concentrations that vary in time and space, microbial population sizes, the effects of immune system responses, and interactions within microbial communities^20–24^. Quantifying the evolution of AMR in clinically relevant settings can provide new opportunities for combatting AMR by informing antibiotic stewardship and developing personalized antibiotic treatments.

Fluoroquinolones are a class of potent broad-spectrum antibiotics that make up a fifth of global antibiotic consumption^25^. In contrast to the majority of antibiotics for which horizontal gene transfer plays an important role in the evolution of resistance^26,27^, fluoroquinolone resistance (FQR) primarily involves mutations in the drug targets (Type II topoisomerases), with mobilized genes playing only a supporting role^28,29^. Analysis of pathogen genomes and *in vitro* experiments suggest that FQR evolves through a stepwise process that begins with specific mutations in DNA gyrase and topoisomerase IV^15,30–32^. However, the specific conditions under which these mutational processes occur in real-world settings remain poorly understood.

Commensal evolution in the gut of adults is evident over a timeframe of several years^33–36^, and over shorter timeframes during the use of antibiotics^37^, after fecal microbiota transplantation^38^, or in newborns^39^. Fluoroquinolone concentrations in the human gut are high and persist days after oral treatment^40^, exerting a strong selective pressure^41^, and disrupting gut communities^42^. We therefore hypothesized that gut commensals would evolve resistance during a single, brief course of ciprofloxacin, allowing us to quantify key aspects of the evolution of AMR *in vivo*.

From a technical perspective, studying *in vivo* evolution in natural settings is challenging. The microbial communities within a single human gut are composed of hundreds of localized populations, each founded through one or several rounds of colonization^35,43^. Gene duplications and horizontal gene transfer result in a convoluted genomic landscape riddled with sequence homology between genomes, resulting in non-trivial relationships between genomes and genome assemblies^36^. One powerful genotyping approach is to sample a community repeatedly over time, which permits sequences to be associated and assembled based on their co-abundance across samples. Repeated sampling has been used to resolve genomes^44–46^ and microbial strains^37,47–49^. However, existing strain recovery tools utilize heuristic approaches to identify true polymorphic sites, which are embedded in a background of spurious variant sites that arise during genome assembly due to sequence homology within and between genomes. As of now, there is no method that effectively leverages dense longitudinal sampling to assemble genomes and accurately identify true polymorphic sites.

In this study, we exposed 60 subjects to a fluoroquinolone antibiotic, ciprofloxacin, orally for five days and collected longitudinal stool samples before and after exposure for a period of 20 weeks to study how gut commensals evolve under antibiotic selective pressure *in vivo*. To identify underlying polymorphisms, we developed a tool (PolyPanner) that simultaneously reconstructed 5665 genomes of localized populations of species and their 2.27M polymorphic variants. PolyPanner is unique in its ability to leverage the statistical power of longitudinal shotgun metagenomic data to address assembly errors and enrich for true polymorphic sites. Analysis of variants that swept in an antibiotic-associated manner uncovered evidence of convergent evolution focused on a pair of amino acids within the gene that encodes DNA gyrase subunit A, *gyrA*, alongside evidence of dispersed selective pressure elsewhere. Nearly 10% of susceptible microbial populations underwent sweeps in gyrA. The probability of a selective sweep was strongly influenced by population abundances both before and during antibiotic exposure. Finally, selective sweeps in *gyrA* and other genes persisted for months after the antibiotic exposure. This study provides valuable insights into the relationship between antibiotic usage and the evolution of resistance in human associated microbiotas.

## RESULTS

Sixty healthy adult volunteers took a short course of ciprofloxacin (500mg orally, twice daily, 5 days) and self-collected 16 stool samples over a period of 20 weeks including daily sampling before, during and after the antibiotic exposure (960 samples in total, see **Supp. Fig. S1**). Stool-derived DNA libraries from all 960 samples were sequenced with 18.8M (million) reads per sample on average. We processed the temporal shotgun reads to generate metagenome-assembled genomes (MAGs) and their polymorphic sites using a workflow that combines standard metagenomic tools and PolyPanner, a program developed here and described in **Methods** (see **Supp. Fig. S2** for an overview of the workflow). Briefly, reads were pooled within subjects to generate 60 subject-specific metagenome co-assemblies spanning ∼400Mb each on average. For each sample, reads were aligned to their respective subject assembly, generating single-nucleotide coverage profiles which represent fluctuations in the abundance of all assembled base pairs over the 16 time points. Chimeric contigs were segmented by applying an algorithm that located abrupt transitions in coverage profiles along contigs (**Methods**). These transitions are likely chimeric assembly breakpoints where the two sides of the breakpoint represent different populations. When applied to our dataset, the algorithm identified ∼10k such breakpoints (involving 2.1% of contigs) per subject on average. Contigs were segmented on these breakpoints and the resulting segments were binned (i.e., clustered) based on average segment-level coverage profiles, resulting in 109k genome bins per subject on average. Downstream analysis was limited to 5665 MAGs (51-166 genomes per subject) that were at least 50% complete and not more than 10% contaminated. On average, the selected genomes were 81.3% complete and 1.8% contaminated (**Supp. Figure S3**), with median sequencing depth (summed across the 16 samples per subject) of 51x (5x to 3856x). Each genome putatively represented the population of a single bacterial species inhabiting the gut of a specific subject. The 5655 populations spanned 924 distinct species, 378 genera and 81 families (**Supp. Table S1**).

### Populations were disturbed yet mostly resilient to the antibiotic exposure

We examined the population-level response to the antibiotic exposure, as reflected by changes in population abundance over time. Population response trends included transient decreases and increases in abundance, indicating relative ciprofloxacin sensitivity and resistance or tolerance, as well as putative extinction events (**Fig. 1A**). During the height of the disturbance (days 3 to 8 after the start of ciprofloxacin) 15.9% of populations were below the detection limit and by the last sampling day (day 77) 9.7% of populations remained undetected and putatively extinct (**Fig. 1B**). Conversely, 0.9% of the populations were not observed at baseline and emerged during the height of the disturbance. The rank order of persistent populations was disrupted during the disturbance (Spearman rho=0.31) and returned closer to baseline by day 28 (rho=0.62). The relatively few extinction events and the overall return to baseline rank of persisting populations attested to the resilience of the human gut microbiome. Response trends were broadly associated with taxonomic identity (**Supp. Fig. S4**), yet diverse taxa such as the *Lachnospiraceae* family were highly heterogenous in their response (**Supp. Fig. S5**). The resilience of gut populations, despite the antibiotic-associated disruption, created a unique opportunity to assess how ciprofloxacin drives evolution in gut commensals.

**Fig. 1.**
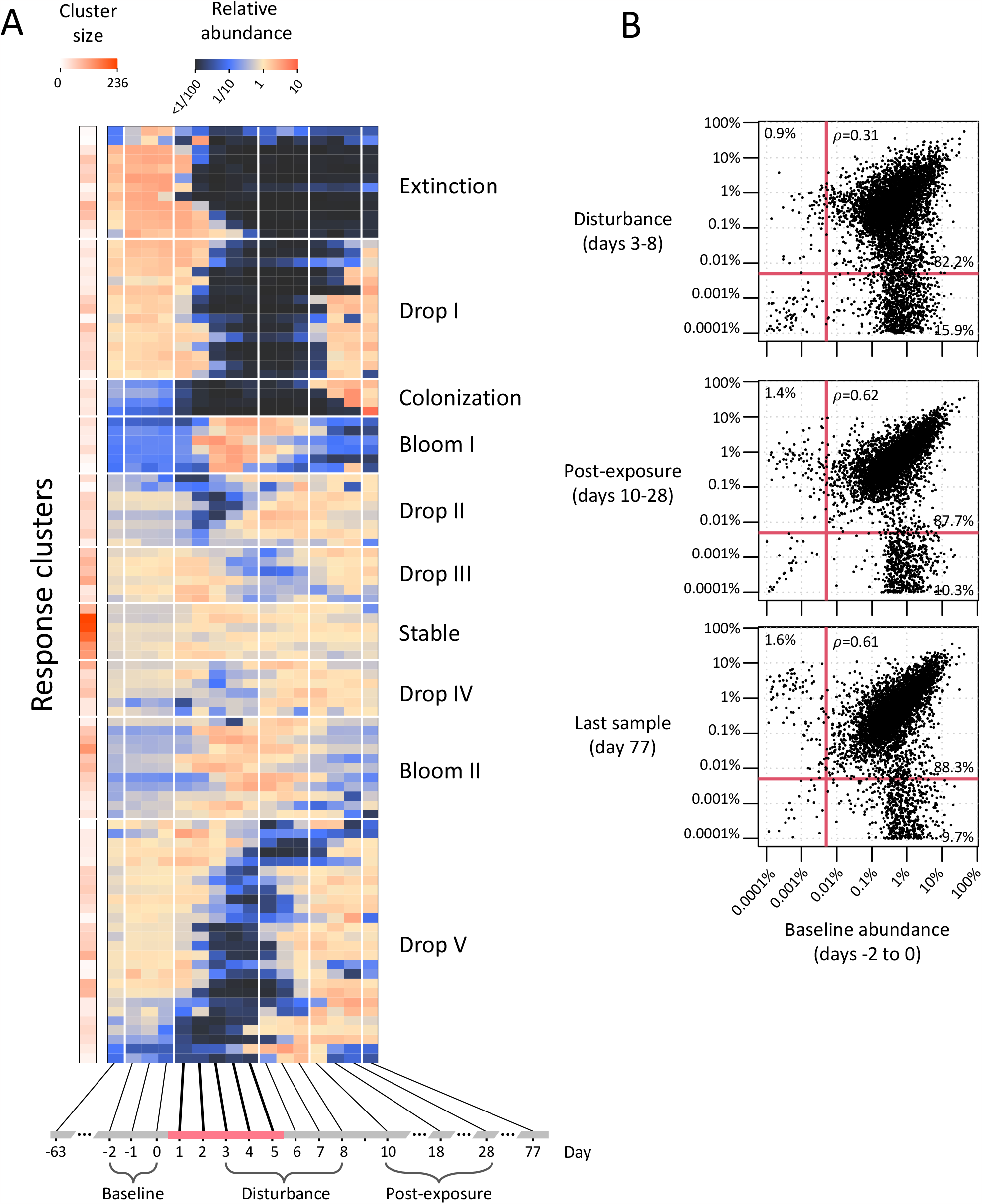
Population-level responses to ciprofloxacin. **A)** Abundance trajectories of all 5665 metagenome-assembled genomes (MAGs) were each normalized relative to their respective mean and clustered using k-means into 100 response clusters or trends. Trends are organized vertically using hierarchical clustering, with the number of populations per cluster shown by a color bar on the left. The mean-centered abundance of each cluster across 16 temporal samples is shown horizontally, with ciprofloxacin course days indicated in red along the axis (shifted by a day to reflect transit time). **B)** Scatter plots comparing the baseline pre-exposure population abundances to three later timeframes. Each scatter plot depicts for each population the baseline abundance (x-axis, average of days -2 to 0) vs. one of three timeframes (during the height of disturbance at days 3-8, post exposure at days 10-28, and on the last day of sampling, day 77). A pseudo-count of 0.0001% was added to each count for visualization purposes. Vertical/horizontal red lines represent a detection threshold of 0.005%. The percentages of populations within quadrants are shown, as are Spearman correlation coefficients between populations detected in both timeframes.

### Identifying polymorphic variants in mixed populations

We wished to identify true polymorphic sites, defined as variant sites that segregated within a single population. Mismatches in read alignments were identified on a single-nucleotide basis, resulting in variant sites typed as single nucleotide polymorphisms (SNPs), short insertions or deletions (indels), and genome rearrangement breakpoints. Sequencing errors were identified using a probabilistic approach and removed (**Methods**), resulting in 107.1 million variants. Due to genomic sequences shared between populations and duplicated genes within populations, not all variants inferred in this manner reflected true polymorphic variants (see illustration of concepts in **Supp. Fig. S6**). We developed a statistical approach to enrich for what we call *dynamic variants*, which are polymorphic variants with a non-constant frequency over time (**Methods**). The remaining variants, which we call *spurious variants*, reflect either homology between distinct populations (*ortholog variants*) or homology within a population due to duplicated genes (*paralog variants*). Spurious variants are considered noise for our purpose of tracking competing alleles within a single bacterial population. To our knowledge, this approach is the first explicit modelling and classification of variant types and allows one to enrich for true polymorphic variants using longitudinal shotgun data.

We assessed the density of spurious variants and benchmarked our ability to remove them by simulating data from 100 bacterial communities with different numbers of introduced dynamic variants (40G reads in total). In these simulated datasets, spurious variants were pervasive, with typically 50-150 spurious variants per Mb of genome, even when few mutations were simulated (**Supp. Fig. S7A**). With our approach, the fraction of spurious variants that were misclassified as dynamic was less than 0.3%, regardless of sequencing depth, meaning that our approach depleted the density of spurious variants by over 100-fold. Sensitivity depended on the sequencing depth and the mutation type. Local mutations (SNPs and indels) began to be detected at a sequencing coverage threshold of 50x per genome and plateaued at 60-80% of variants detected (failing to reach 100% due to incomplete genome assemblies) when coverage was above 100x (**Supp. Fig. S7B**). The sensitivity of detection for large-scale genome rearrangements was lower (**Supp. Fig. S7C**). Importantly, the precision remained high in the face of “genetic crowding” in which communities contained multiple members of the same genus or family (**Supp. Fig. S8**). Taken together, simulated data demonstrated the pervasiveness of spurious variants and showed that our approach is well-suited to detect dynamic variants in evolving bacterial communities.

### Hundreds of populations underwent putative selective sweeps

Application of our approach to the 5665 populations detected 2.27 million dynamic variants, making up 2.1% of all variants (“Genome” table in **Supp. Table S1**). Dynamic variants were comprised of 98.5% SNPs, 1.3% indels and 0.2% rearrangement breakpoints. Dynamic variants were found in 2133 populations (38%), and the percentage of populations with dynamic variants among sufficiently sequenced populations (with an abundance above ∼1%) was 85.8% (**Fig. 2A**). We limited the analysis to 1866 populations that had 1-1000 dynamic variants, omitting 267 populations with over 1000 variants that were composed of distant conspecific strains. Underlying strain genomes and their temporal trajectories were approximated using a probabilistic model^48^, resulting in 2-8 strains per population (**Fig. 2B**).

**Fig. 2.**
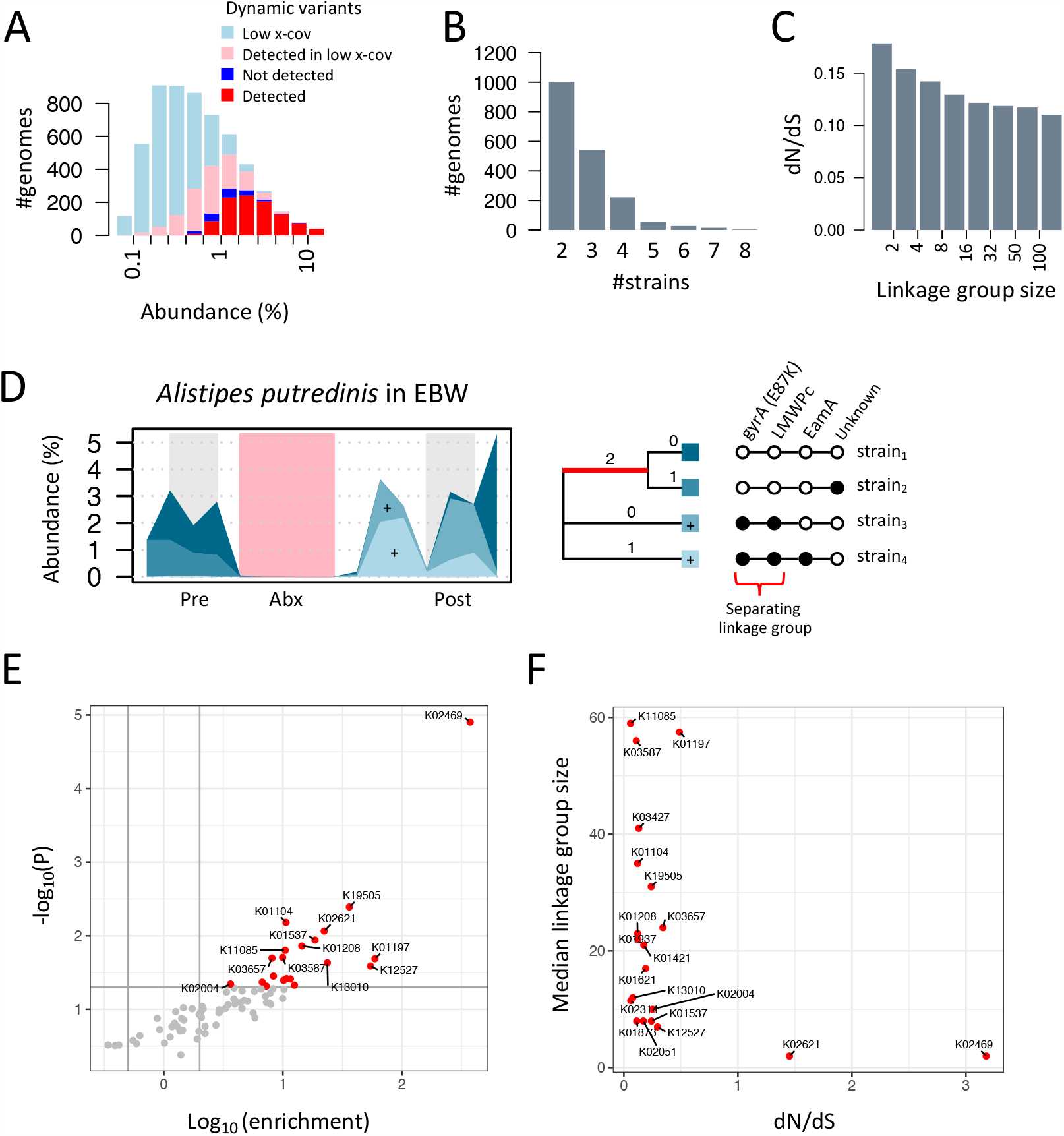
Variants, strains, linkage groups and converging evolution analysis. **A)** Number of population genomes (y-axis) as a function of the average population abundance (x-axis). Colors denote four population types: 1018 populations with dynamic variants and sufficient x-coverage to detect sweeps if present (red, “Detected”), 168 populations with no dynamic variants and sufficient x-coverage (dark blue, “Not detected”), 1115 populations with dynamic variants and low x-coverage (pink, “Detected in low x-cov”), and 3364 populations with no variants and low x-coverage (light blue, “Low x-cov”). **B)** Number of genomes containing dynamic variants (y-axis) as a function of the number of inferred strains (x-axis). **C)** dN/dS ratios (y-axis) as a function of the linkage group (LG) size (x-axis). **D)** Temporal dynamics of the population of *Alistipes putredinis* in subject EBW. On the right is the variant-strain matrix, with 4 bi-allelic polymorphic loci (columns, named according to gene), and 4 strains (rows), with black and white circles depicting the variants present in a strain. The strain phylogeny (generated through maximum parsimony) is shown in the middle, with the number of switching variants displayed above each edge. The edge associated with the linkage group that separates strains 1+2 from strains 3+4 is colored in red. Shown on the left is the temporal abundance of strains (strains stacked) over the 16 temporal samples (x-axis), with background colors emphasizing pre (antibiotic) samples (days -2 to 0, gray), disturbed samples (days 3-8, pink) and post (antibiotic) samples (days 10-28, gray). Plus signs denote strains 3 and 4, which become the dominant strains after the exposure. Strain order and representation as shades of blue are consistent across the panel. **E)** Variants were assigned weights equal to the inverse of their smallest linkage group, weights were aggregated within KEGG Orthology annotations (KOs) of associated genes and then compared to a null model of weights across KOs generated through permutations. A volcano plot depicts weight enrichment ratios over the null model (x-axis) vs. P-values (y-axis). Significant KOs (P<0.05) are colored red. K02469 is *gyrA* and K02621 is *parC*, other KO descriptions are given in **Supp. Figure S12. F)** For each significant KO, the dN/dS ratio (x-axis) of the supporting variants is plotted vs. the median linkage group size of the supporting variants (y-axis).

For each population, strains were arranged on a most-parsimonious phylogenetic tree, and each tree branch was associated with a *linkage group* (LG), defined as the set of variants that were inferred to change along the branch. We considered the LG size a crude estimator of the mean age of the variants in the linkage group, where small linkage groups indicated recent events (including *de novo* mutations and recombination events), and large groups earlier events. Smaller LGs were correlated with elevated levels of non-synonymous mutations, measured using dN/dS ratios (**Fig. 2C**).

There were 7132 variants that “swept” from a pre-antibiotic frequency below 20% to over 80% post-antibiotic (“Variant” table in **Supp. Table S2**). Each sweeping variant represented a putative mutation that transitioned from the baseline to the post-exposure variant state. Sweeping variants involved 513 populations that were distributed across all subjects (8.55±4.15 populations per subject) and involved 212 species (22.9% of species). Sweeping variants rarely traveled alone: only 1% were the only member in their LG and 6.5% had 2-10 variants in their LG (**Supp. Fig. S9**). We illustrate how variants are organized into strains and linkage groups using a *Alistipes putredinis* population found in subject EBW, where an antibiotic-associated shift in strain composition could be attributed to an LG composed of 2 mutations, one of which was in the gyrase subunit A (*gyrA*) (**Fig. 2D**).

### *Convergent evolution focused on* gyrA

A strain can sweep following antibiotic exposure because it carries an adaptive ‘driver’ mutation, but additional ‘passenger’ variants in the same strain may also be swept to high abundance^50^. We therefore searched for evidence of evolution occurring in parallel in different populations and subjects. To reduce the background noise of passenger variants, we focused our analysis further on 1771 sweeping variants (in 312 populations) that were part of small LGs (up to 100 variants per LG). Variants were associated with single genes if intragenic or with two adjacent genes if intergenic. Interestingly, 22.5% of implicated genes were associated with multiple variants in the same population, indicating that at least some of the genetic variation was likely introduced through recombination (**Supp. Fig. S10**).

Genes were annotated using the KEGG Orthology (KO) database, with special attention to distinguishing between the homolog genes, gyrase subunit A (*gyrA*, K02469) and topoisomerase IV subunit A (*parC*, K02621)^51,52^ (**Supp. Fig. S11** and **Supp. Table S3**). Variants were assigned a weight inverse to the size of their LG, and weights were aggregated within KOs, resulting in 803 KOs with positive weights. Weight enrichment ratios were computed over a background model in which variants were randomly reassigned to a gene within their genome, resulting in 20 significant KOs that showed evidence of convergent evolution (**Supp. Fig. S12** and **Supp. Table S4**). The top hit in terms of fold-enrichment and statistical significance was *gyrA* (68 variants, 373-fold enrichment, *P*< 1.25 × 10^!”^) (**Fig. 2E**). *GyrA* sweeps occurred in 63 populations inhabiting 34 subjects. A dN/dS ratio of 3.9 and a median LG size of 2 were highly suggestive that *gyrA* mutations occurred *de novo* within the gut of the subjects (**Fig. 2F**). See **Supp. Fig. S13** for examples of *gyrA*-associated sweeps.

Beyond *gyrA*, the selective pressure was less focused and was distributed among the remaining 19 KOs, with no single KO involving more than 6 populations. Breakdown of KOs by taxonomic identity showed how *gyrA* sweeps were widely distributed (41 species, 15 families), while other KOs were typically limited to specific taxa (**Supp. Fig. S14**). Notably, *parC* was enriched in mutations yet to a lesser degree (only 5 variants in 3 families) than its close homolog *gyrA*, possibly because of reduced selective pressure due to its distinct role compared to DNA gyrase^53^, and in contrast to previous findings suggesting that Gram positive bacteria evolve primarily through mutations in *parC*^28^. Other implicated KOs included efflux pumps (K02004), DNA helicases (K02314, K03657), transport systems (K01537, K02051, K11085), transcription factors (K19505) and phage-associated genes (K01421). The inhibition of the helicase DnaB (K02314) has been implicated previously in FQR in *E. coli*^54^. While parallel evolution beyond *gyrA* was limited, there were compelling examples of idiosyncratic sweeps that were not driven by *gyrA* (**Supp. Fig. S15**).

### Selective pressure targets specific amino acids within gyrA

The mutations associated with *gyrA* were concentrated at two residues: *gyrA*:83 (56 substitutions) and *gyrA*:87 (9 substitutions) (*E. coli* numbering) (**Fig. 3A**). Substitutions at these two positions are well-documented as initial steps towards fluoroquinolone resistance^29^, and accounted for 87% of *gyrA* substitutions observed in our study. We focused on *gyrA*:83 substitutions, which was by far the most common in our study, occurring in 33 subjects and in 38 species that spanned 11 taxonomic families. *GyrA*:83 substitutions were mainly serine to phenylalanine (S83F, 21 cases) and serine to leucine (S83L, 19 cases) (**Fig. 3B**). To assess substitution probabilities at position 83, we first resolved *gyrA*:83 at baseline for 3896 genomes that contained *gyrA*. Serine, which is known to be associated with fluoroquinolone susceptibility^15^, was present at baseline in 61% of genomes, while leucine and phenylalanine, associated with resistance, were present in 6% of the genomes (**Fig. 3C**). Baseline amino acids at *gyrA*:83 were associated with taxonomic identity (**Supp. Fig. S16**). The probability of serine substitution was 9.6±1.5% per population (**Fig. 3D**). In other words, nearly 10% of susceptible bacterial populations evolved towards resistance *in vivo*. To our knowledge this is the first estimation of the rate at which ciprofloxacin resistance evolves in human subjects.

**Fig. 3.**
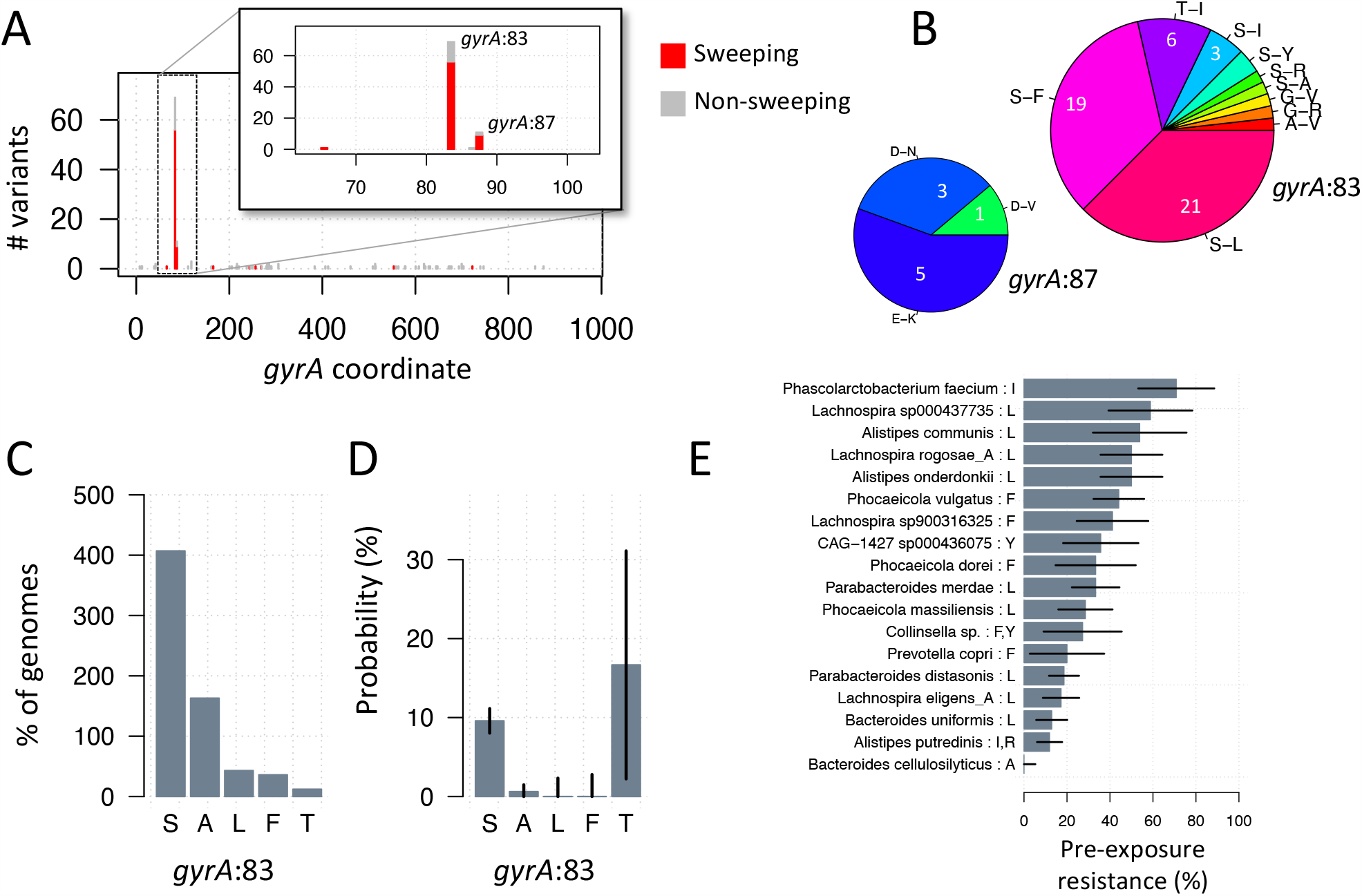
Focused selective pressure on *gyrA*. **A)** The number of variants (y-axis) for each amino acid (AA) position within the *gyrA* gene (x-axis, *E. coli* numbering). Sweeping and non-sweeping variant counts are colored red and gray respectively. **B)** Breakdown of substitutions associated with sweeping variants at *gyrA*:83 and *gyrA*:87, letters represent amino acids (e.g., ‘S-F’ represents a substitution of serine by phenylalanine). **C)** Breakdown of all genomes containing *gyrA* according to the baseline AA identity at *gyrA*:83 before the exposure. **D)** Exposure-associated substitution probability, stratified by the baseline AA at *gyrA*:83. Confidence intervals showing the standard deviation depicted with vertical lines. **E)** Species-specific resistance alleles at *gyrA*:83 were determined for 31 species based on observed exposure-associated substitutions. The percentage of resistance alleles prior to exposure is shown for 18 species represented by at least 10 populations. Resistance alleles are shown next to the species name. For example, *Alistipes putredinis* has two resistance alleles at *gyrA*:83, namely isoleucine (‘I’) and arginine (‘R’), and 12% (5/42) of *A. putredinis* populations had either of these alleles prior to the exposure. Black bars represent standard deviation estimates.

Resistance mutations at *gyrA*:87 appeared less favorable. There were only 9 such instances, and *Alistipes putredinis* accounted for five E87K mutations that likely occurred because these genomes have an unusual initial serine codon at *gyrA*:83 (AGU) that would require two mutations to reach the more favorable leucine or phenylalanine at position 83. Clinical isolates of *Neisseria gonorrhoeae* were recently shown to gain resistance through mutations in *gyrA*:87, while keeping serine at *gyrA*:83^55^.

### Evidence for pre-existing resistance

We wished to leverage the large scope of the dataset to assess the prevalence of FQR prior to ciprofloxacin exposure in our subjects. We determined species-specific resistance alleles at *gyrA*:83 for 38 species, based on the identity of the substituting amino acid in sweeping variants. Remarkably, resistance alleles were observed prior to the ciprofloxacin exposure in 26 of the 38 species (**Supp. Fig. S17**). For example, the resistance allele of *Phocaeicola vulgatus* was phenylalanine (based on 2 sweeps with S83F), and there were 15 out of 34 populations with phenylalanine at *gyrA*:83 before the exposure. Frequencies of resistant strains at baseline varied among species, ranging between 0% to 71% and averaging 33.8% (**Fig. 3E**). Subjects did not have significant differences in the overall level of pre-existing resistance, suggesting that most strains were likely resistant prior to colonization, rather than evolving resistance *in situ* within subjects (**Supp. Fig. S18**). Some species had a signature of pre-exposure resistance (based on phenylalanine or leucine at *gyrA*:83), despite not having any observed sweeps in our study. For example, the percentage of resistant *E. coli* strains was estimated to be 41% at baseline (based on 5 leucines and 7 serines at *gyrA*:83), without any observed *gyrA* sweeps in *E. coli* populations.

### Evolvability of resistance predicted by population size and drug sensitivity

An important question with possible clinical implications is what determines the evolvability of a population when challenged with ciprofloxacin, which is the probability that a population will have one or more sweeping variants. For this analysis we focused on 410 populations (out of 5665) that had serine at *gyrA*:83 at baseline and had sufficient coverage for detection of sweeps, if present (equivalent to an abundance over ∼1%, as shown in Figure 2A). The overall evolvability was 0.26 based on 106 populations with one or more sweep. *GyrA* evolvability (where at least one sweep involves *gyrA*) was 0.11, based on 45 populations. Thus, *gyrA* evolution accounted for only 36% of the overall evolvability, highlighting the putative role of alternative evolutionary trajectories in response to antibiotics.

We employed logistic regression to assess whether abundance and taxonomic identity can serve as reliable predictors for the *gyrA* evolvability of a population (**Sup. Figure S19**). Logistic regression identified four significant predictors: baseline population abundance (P<0.02), the decline in abundance during antibiotics (average of days 1-5, P<10^-5^), and phylum (Bacteroidetes P<2x10^-3^and Actinobacteria P<2x10^-8^, relative to Firmicutes) (**Fig. 4A**). In terms of effect sizes, *gyrA* evolvability exhibited a 3.8-fold increase for every 10-fold rise in baseline population abundance and a 3.2-fold increase for every 10-fold rise in the magnitude of decline in abundance during antibiotic exposure (i.e., the ratio between abundance at baseline and during treatment). *GyrA* evolvability showed a strong association with drug sensitivity, as 90.2% of the populations that underwent a *gyrA* sweep experienced a reduction in abundance during antibiotic exposure. *GyrA* evolvability of Bacteroidetes and Actinobacteria was 3.8-fold and 26-fold higher than Firmicutes, respectively. For the most susceptible populations (i.e., with a decline of 1000-fold or more during antibiotics), evolvability equaled 0.5 when the baseline abundance was ∼0.1% for Bacteroides and ∼1% for Firmicutes. Model-predicted evolvability and observed evolvability were in agreement, when stratifying populations by baseline abundances (**Fig. 4B**), or by the decline in abundance during antibiotics (**Fig. 4C**).

**Fig. 4.**
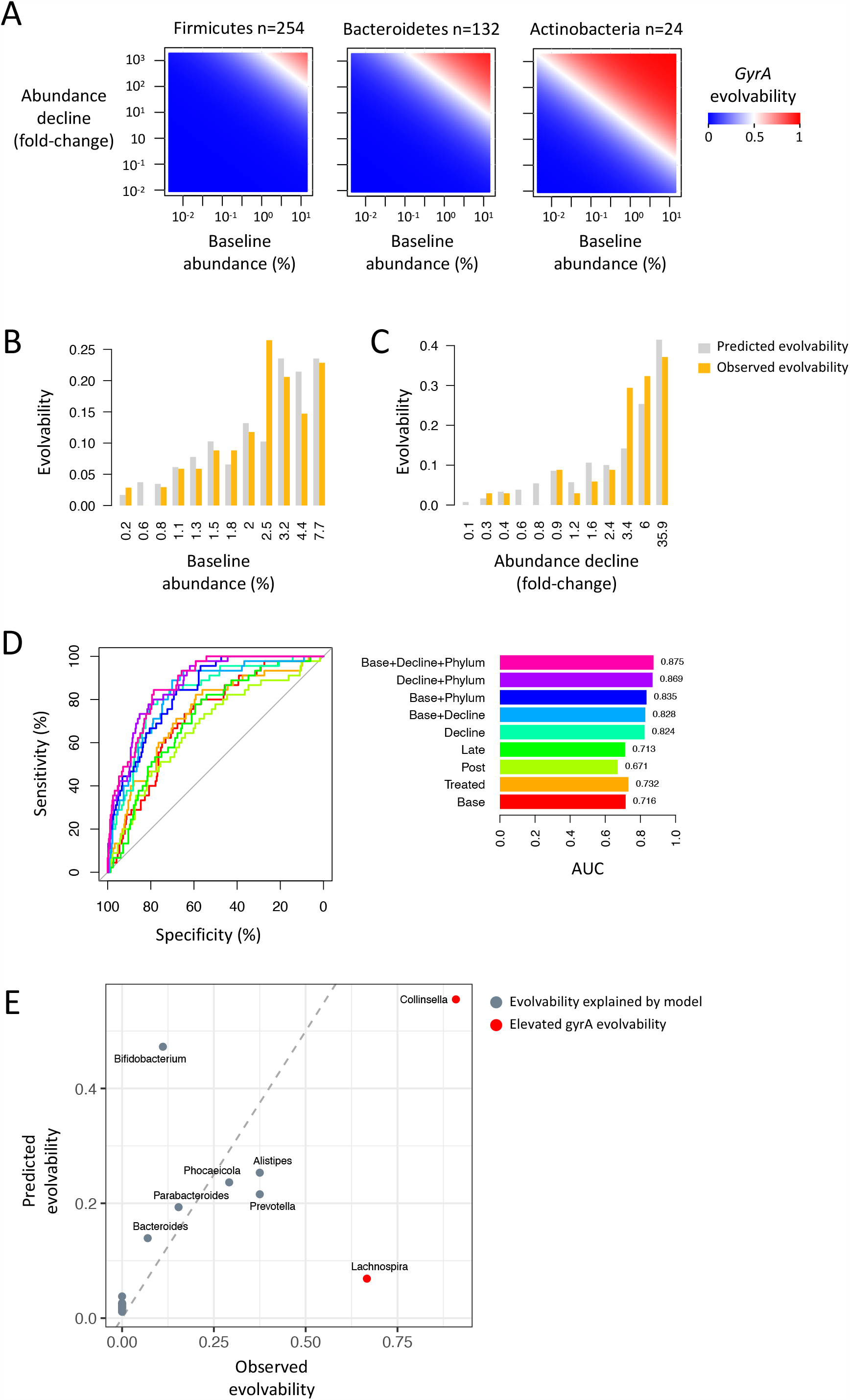
Abundance-dependent evolution of antibiotic resistance. **A)** *GyrA* evolvability as predicted by a fitted logistic regression model based on variables for the baseline population abundance (average of days -2 to 0, x-axis), the fold-change decline during antibiotics (average of days 1-5, y-axis), and phylum (separate plot for each of three phyla). **B)** Model-predicted and observed *gyrA* evolvability (y-axis) as a function of baseline abundances. **C)** Model-predicted and observed *gyrA* evolvability (y-axis) as a function of the fold-change decline in population abundances during antibiotics (days 1-5). **D)** Shown on left are ROC curves (receiver operating characteristic curves) of 9 inferred logistic regression models that predict the probability of a population to have a sweeping *gyrA* variant. Shown on right are the areas under the curve (AUC) of all models. **E)** Observed (x-axis) vs. model-predicted (y-axis) genus-level *gyrA* evolvability, for the 21 genera associated with at least 6 populations. Genera are colored red if differences between observed and predicted evolvability were significant (Binomial distribution, P<0.05), and colored gray otherwise. Dashed line depicts the line x=y.

The model performance was excellent, with an area under the curve (AUC) of 0.875 (**Fig. 4D**). Of note, a simple model that solely relies on the baseline abundance had an AUC of 0.76 and is distinctive in its capability to predict evolvability using pre-exposure data. While differences in genus-level *gyrA* evolvability were captured by the model in general, *Collinsella* (phylum Actinobacteria) and *Lachnospira* (Firmicutes) were significantly more evolvable than predicted by the model (**Fig. 4E**). Non-*gyrA* evolvability (i.e., the probability of a sweep with a gene other than *gyrA*) was only partially explained by the abundance during antibiotics but not by the abundance at baseline and was significantly harder to predict, reaching a maximal AUC of only 0.567 (**Supp. Fig. S20**). Evolvability was not significantly different between subjects (**Supp. Fig. S21**).

The model parameters of *gyrA* evolvability are related to fundamental attributes of bacterial populations: baseline abundances are linked to absolute population sizes and the magnitude of the decline during antibiotics is a crude proxy for sensitivity to ciprofloxacin. The data and models are consistent with an enrichment of *de novo gyrA* mutations that swept primarily in populations that were both abundant and sensitive through a bottleneck event, while evolution through non-*gyrA* variants had no clear signature of *de novo* mutations, likely drew from pre-existing standing variation, and was not clearly associated with resistance.

### Sweep reversion was slow

Finally, we wanted to determine the long-term effect of the exposure, by examining sweep reversions. At the time of the last sample, collected 10 weeks after the end of the exposure, 89% of *gyrA* sweeps (65% of non-*gyrA* sweeps) remained at a frequency above 80% (**Fig. 5A**), and 74% of *gyrA* sweeps (56% of non-*gyrA* sweeps) were at a frequency of 100% (**Fig. 5B**). To predict the behavior beyond 10 weeks, we employed a naïve model that assumed a constant fitness cost for resistant alleles, and separately inferred selection coefficients for all sweeps; see an example of the fastest reverting *gyrA* sweep in **Fig. 5C**. Projecting one year out, 95% of *gyrA* sweeps were not expected to fall below 1% frequency, which was higher than 84% for non-*gyrA* sweeps (P<0.029, Yates’s chi-squared test). The median reversion time for sweeps that were projected to revert within a year was 109 days (**Fig. 5D**). Taken together, our data indicate that mutations in *gyrA* and other loci can persist long after the antibiotic exposure, consistent with relatively low fitness costs or compensatory mutations.

**Fig. 5.**
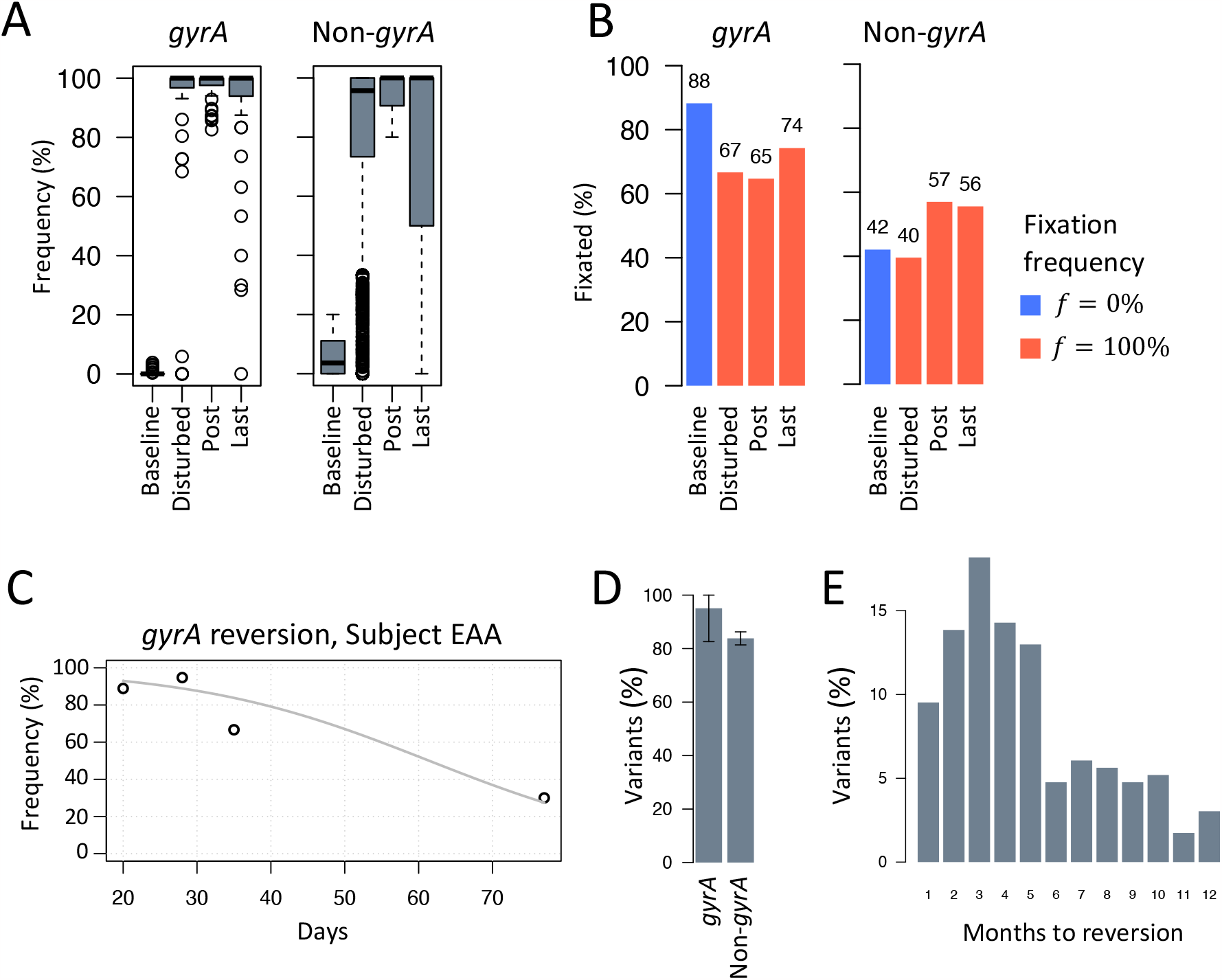
Long-term persistence of antibiotic-associated genetic changes. **A)** Boxplots depicting the distribution of the frequencies of sweeping *gyrA* variants (left) and non-*gyrA* variants (right), stratified by time windows (day ranges defined in panel 1A). Gray rectangles span the first and third quartiles, horizontal line depicts the median, dashed line spans between the 5^th^ and the 95th percentiles, and outliers outside that range are plotted as circles. **B)** The percentage of sweeping variants fixated at 0% frequency at baseline (blue) or fixated at 100% within the 3 other time windows (red), shown separately for *gyrA* variants (left) and non-*gyrA* variants (right). **C)** Example of a sweeping *gyrA* variant in subject EAA that was projected to revert below a frequency of 1% within 135 days. Circles depict the variant frequency (y-axis) as a function of the day relative to the start of the antibiotics (x-axis). Gray line shows an exponential decay, based on a constant selection coefficient of 0.006 that was inferred from the temporal frequency data, assuming 10 generations per day. **D)** The percentage of sweeping *gyrA* variants (left) and sweeping non-*gyrA* variants (right) that were projected to require more than a year to revert below a frequency of 1%. Vertical lines depict estimates of the standard deviation. **E)** For sweeping variants that were projected to fall below a frequency of 1% within a year, a histogram of the number of months until reversion.

## DISCUSSION

This work describes how gut commensals evolve *in vivo* during and after a single short course of ciprofloxacin. We quantified, for the first time, rates at which AMR and other antibiotic-associated mutations sweep and persist in human gut communities under clinically relevant conditions. Our analysis shows that 11% of commensal populations acquire putative resistance through mutations in *gyrA*, and over 80% of acquired genetic changes remain dominant months after the antibiotic exposure. Evolvability through *gyrA* was explained by a simple model based on population abundances, with marked differences between phyla. The study reveals key factors that affect the evolution of ciprofloxacin resistance in humans.

Our data show that *gyrA* evolvability of bacterial populations surpasses a probability of 0.5 when population abundances cross a threshold of 0.1% and 1%, for Bacteroides and Firmicutes respectively, which together make up 86% of populations. Yet conservative assumptions result in a theoretical upper bound for this threshold that ranges from 0.0002% to 0.002% (**Supp. Note 1**). The large discrepancy between theory and empirical data, and between phyla, may reflect differences in mutation rates and sweep frequencies, and can also be explained by reduced effective population sizes in the intestines due to flow^56^. Recent *in vitro* work with *E. coli*^15^ and *Pseudomonas aeruginosa*^16^ showed that *gyrA* mutations sweep only when populations are large, in agreement with our *in vivo* results. The emerging picture is that large populations can evolve mutations in *gyrA* on their evolutionary trajectory towards higher levels of fluoroquinolone resistance, while smaller populations may evolve through a variety of other mutations, if at all, and are more likely to lose their resistance over time. From a clinical perspective, precise models of evolvability can inform the selection of appropriate antibiotics, as previously suggested from *in vitro* work^32^. To give a concrete example, a clinician treating a patient with a urinary tract infection may decide against prescribing fluoroquinolones if sequencing data suggest that the abundance of a gut-residing pathogen is high enough to predict evolution of resistance during treatment. In the future, routine assessments of microbial composition and abundance could become a useful tool for mitigating risk for the patient and society^3^.

The method presented here exploits temporal shotgun metagenomic data, allowing us to re-examine decades of *in vitro* studies and provide a more relevant understanding of how AMR evolution transpires *in vivo*. The approach is well-suited to detect adaptive evolution in diverse natural microbial communities facing selective pressure, although it has some limitations. Studying horizontal gene transfer necessitates complementary methods like Hi-C^36,57^ and long-reads^37,58–60^. Additionally, the method is limited in its ability to detect ‘soft’ sweeps in which multiple independent mutations sweep simultaneously^61^. Such phenomena, supported by *in vitro* data, could play a significant role in the evolution of antimicrobial resistance^15^.

The persistence of *gyrA* mutations may suggest that they have little or no fitness cost, a well-known concern^28,62^. For example, *in vitro* experiments showed that replacing serine at *gyrA*:83 had no measurable fitness cost in *E. coli* and *Streptococcus pneumoniae*^63^, or even conferred a fitness advantage in pathogens such as *Salmonella* Typhi and *Campylobacter jejuni*^64,65^. For mutations to revert, their fitness cost must supersede the nominal selective pressure in the gut for a substantial length of time, yet the breadth of selective pressures under natural conditions (other than antibiotics) remains poorly understood.

This approach can be used to understand the evolution of resistance *in vivo* with different antibiotic classes, dosages, and durations, as well as previously unrecognized effects of other drugs and conditions that impose selective pressure on the microbiome. The results of such studies may inform the design of revised antibiotic regimens that better optimize benefit/cost ratios^66,23^. By studying microbial communities in their natural environment, the net effect of myriad interacting externalities on the evolution of community members can be distilled and eventually understood.

## ACKNOWLEDGMENTS

We extend our gratitude to Benjamin Good and Elizabeth Costello (Stanford University) for their valuable insights and constructive feedback. This work was funded by the National Institute of Allergy and Infectious Diseases of the National Institutes of Health grants R01AI147023 (DAR) and R01AI112401 (DAR), and the Thomas C. and Joan M. Merigan Endowment at Stanford University (DAR).

## AUTHOR CONTRIBUTIONS

Conceptualization: EY, LD, DAR; Data Curation: EY, LD; Formal Analysis: EY; Investigation: AVP, CG; Methodology: EY, SH; Software: EY; Supervision: EY, DAR; Validation: EY; Visualization: EY; Writing – Original Draft Preparation: EY; Writing – Review & Editing: EY, LD, DAR.

## DATA AVAILABILITY

The source code for PolyPanner is available at https://github.com/eitanyaffe/PolyPanner. Unprocessed DNA sequence reads are available in the NCBI database, under project PRJNA974858.

