## Supplementary material for "A short course of antibiotics selects for persistent resistance in the human gut": Methods and Supp. Notes

### Human subjects, sample collection, basic processing

**Sample collection and DNA sequencing.** This study was approved by the Institutional Review Board of Stanford University (protocol #25268). All participants provided written informed consent before completing an enrollment questionnaire and providing biological samples. 60 heathy adults living in the U.S. took ciprofloxacin for 5 days (500mg orally, twice daily). Each subject self-collected stool samples 9 weeks prior and for each of the 2 consecutive days immediately prior to the start of ciprofloxacin, daily during antibiotic exposure (Days 0-4) and for the following four days (Days 5-8), and then on Days 10, 18, 28, and 77, following the sampling scheme described in Supp. Figure S1. Subjects had not taken any antibiotics for at least 6 months prior to the start of sampling. Samples were kept temporarily at home at -20C, shipped to the laboratory on dry ice and stored at -80C until processed. DNA was extracted using the AllPrep DNA/RNA Mini Kit (Qiagen), sheared and size-selected (>300bp), and DNA libraries were sequenced (2x150nt) at the Chan Zuckerberg Biohub using the NovaSeq 6000 platform.

**Processing raw reads**. Libraries were rarified to 50M read pairs if there were more than 50M read pairs, resulting in 1.9M to 50M read pairs per library (median: 17.46M). Adapter removal and quality filtering were performed using Trimmomatic[67] (v0.38), with the parameters "ILLUMINACLIP:NexteraPE-PE.fa:2:30:10:1:true LEADING:20 TRAILING:3 MAXINFO:60:0.1 -phred33". Duplicate read pairs (identical matches on both sides) were removed. Read pairs that mapped to the human genome were discarded using DeconSeq[68] (v0.4.3, hg38 as reference).

**Metagenome co-assembly and read alignment**. Reads were pooled per subject and co-assembled into subject-specific co-assemblies using MEGAHIT[69] (v1.2.9) with parameters "--min-contig-len 200 --k-min 27 --k-max 77 --k-step 10 --merge-level 20,0.95". Contigs shorter than 1kb were discarded. Read sides were mapped to their co-assembly using BWA-MEM[70] (v0.7.17) resulting in SAM (Sequence Alignment Map format) files. Reads with low-quality alignments (>20 mismatches, <50nt match length or mapping score <30) were removed.

### MAGs and dynamic variants

**Approach overview.** We developed PolyPanner, a program that leverages dense temporal sampling to improve assembly quality and identify high confidence polymorphic variants. PolyPanner receives as input a set of shotgun libraries that are aligned to their co-assembly in SAM format. It transforms the alignments to single-nucleotide coverage vectors that represent library-specific read counts of perfect and mismatch alignments at each base pair in the co-assembly. Tasks performed by PolyPanner are (1) contig refinement; (2) genome trimming; (3) removal of sequencing errors; and (4) identification of dynamic variants.

**Read alignment representation.** For each library, a library data structure was generated from the SAM files and the subject-specific co-assembly as follows. Reads clipped on both sides were discarded, based on having an 'H' or 'S' on start and end of the CIGAR string (Compact Idiosyncratic Gapped Alignment Report). To ensure a margin of safety at the start and end of reads, each read was trimmed by at least 20nt on both sides and trimmed further to avoid a possible overlap between paired reads. Clipped reads were piled-up to generate single-nucleotide coverage vectors that allowed a query as to how many reads covered a query position or a query genomic interval. CIGAR strings were parsed to identify 4 types of variants: (1) a substitution was defined by a source nucleotide (in the contig) and target nucleotide (in the read), as specified by the MD field in the SAM file; (2) a deletion was defined by the number of deleted nucleotides; (3) an insertion was defined by the sequence added between two adjacent positions; and (4) a rearrangement was assigned to the left or right of the position at which the alignment of a clipped read terminated, and was defined by the identity of the contig to which the paired read mapped, if present. Variants were recorded within each library according to their complete identity (position and associated fields). For example, 2 variants representing the insertion of AA and AAA at the same position were counted separately.

**Removing sequencing errors.** Sequencing errors were removed following the rigorous approach taken by Quince et al.[47] We extended their test, which identifies bi-allelic positions that segregate through substitutions, to identify multi-allelic positions that segregate through all types of variants. We inferred 4 global error coefficients $\epsilon_{sub}, \epsilon_{indel}, \epsilon_{rearrange}, \epsilon_{none}$ that represent single nucleotide substitution errors, insertion/deletion errors, rearrangement errors, and no sequencing errors respectively. All libraries of a subject were merged into a single library for this work. Error coefficients (except $\epsilon_{none}$) were seeded at 0.01 and summed to 1. The algorithm repeated the following two steps until convergence: (1) inferring true variants while keeping error coefficients constant, and (2) inferring error coefficients while keeping true variants constant. True variants at a specific position were identified as follows. Let $v_0, v_1 \ldots, v_{n-1}$ denote the variants at the position and their respective read support coverage $t_0, t_1 \ldots, t_{n-1}$, with read supports sorted high to low. Hypothesis $\mathcal{H}_i$ is that variants $v_0, \ldots, v_i$ are true and the remaining variants are a result of sequencing errors. The likelihood of the hypothesis is the multinomial $\mathcal{H}_i(t_0, t_1 \ldots, t_{n-1}|\epsilon) = \frac{T!}{\prod_j t_j!} \prod_j \left(\sum_{k=0,\ldots,i} w_k \times \epsilon_{k,j}\right)^{t_j}$, where $T = \sum_j t_j$, $w_k$ is an approximation of the true frequency of variant $v_k$ that equals 1 if $i = 0, k = 0$ and equals $\frac{t_k}{T}$ otherwise, and $\epsilon_{k,j}$ is the error coefficient that represents a transition from variant $v_k$ to $v_j$. Coefficient $\epsilon_{k,j}$ was determined based on the first condition met: (1) $\epsilon_{k,j} = \epsilon_{none}$ if the variants were identical, (2) $\epsilon_{k,j} = \epsilon_{rearrange}$ if either variant was a rearrangement, (3) $\epsilon_{k,j} = \epsilon_{indel}$ if either variant was an insertion or deletion, and (4) otherwise $\epsilon_{k,j} = \epsilon_{sub}$. For each variant $v_i$ ($i > 0$, $t_i > 2$) we applied the likelihood ratio test $-2log\frac{\mathcal{H}_{i-1}}{\mathcal{H}_i}$, which is approximately distributed as a chi-square distribution, and used the test to assess $p$-values for the hypothesis that a variant is present at a specific position[47]. A Benjamini-Hochberg correction was applied with a false discovery rate (FDR) of 0.001 to account for multiple testing, resulting in a list of true variants. In the second step, error coefficients were inferred while keeping true variants fixed and enforcing a minimal error rate of 0.001% and a maximal error rate of 5%. Each error coefficient was approximated by the average error rate for all positions that do not contain a true variant. The two steps were repeated with variants reclassified and coefficients re-estimated until the set of true variants converged.

**Linkage test.** We denote by $r_i(x)$ the number of reads that cover position $x$ in library $i$ (also called the x-coverage of the position). We denote by $r(x)$ the position coverage vector across libraries $r(x) = (r_1(x), \ldots, r_m(x))$ , where $m$ is the number of libraries. We denote by $r_i(x, y)$ the number of reads fully contained in a sequence interval $[x, y]$ in library $i$, and by $r(x, y)$ the interval coverage vector $r(x, y) = (r_1(x, y), \ldots, r_m(x, y))$. We handle variants at a position in a similar fashion, with $r_i(v)$ denoting the number of reads supporting the variant in library $i$, and with $r(v)$ denoting the variant coverage vector. A pair of sequences (either two positions, two intervals, or two variants) are called *separated* (or *non-linked*) if their associated coverage vectors are independent, based on a Pearson's chi-squared test of independence (applied with a pseudo-count of 0.1 and requiring $P < 0.01$). Note that two sequences for which the associated coverage vectors were not significantly independent are either perfectly linked (i.e., co-occurring in all genomes), or the coverage depth is not high enough to detect separation.

**Co-assembly refinement.** Given a genomic position $p$, we define the left and right intervals $L_p = [p - D, p - d]$ and $L_p = [p + d, p + D]$, where $d = 10$ and initially $D = 200$. The interval $[p - D, p + D]$ is called the spanning interval of position $p$. The position is called a *separating* position if the left and right intervals are separated as defined above. Each co-assembly contig was refined as follows. We tested for separation all positions in the contig associated with a rearrangement variant, and positions distributed across the contig (50bp apart). Separating positions (P<0.01) with a spanning interval entirely contained in the contig were considered candidate breakpoints. The contig was then processed recursively by selecting a single candidate breakpoint at each step. The selected breakpoint was either the candidate breakpoint associated with a rearrangement variant that was supported by the highest number of reads (if such a breakpoint existed), or the candidate breakpoint with the highest chi-square statistic (if no candidate rearrangement breakpoint were found). Only breakpoints with a spanning interval that did not contain any previously selected breakpoints were considered. The contig was split into two segments at the selected candidate breakpoint and the process continued recursively on both segments until no candidate breakpoints were found. After the recursion ended, the induced segments were further refined using the same procedure but with $D = 400$. Finally, a Benjamini-Hochberg correction was applied (FDR of 0.25) to the $p$-values of the breakpoints used to separate the contig into segments, rejecting breaks above that threshold. The result was a final list of breakpoints and the corresponding induced segments.

**Genome binning and trimming.** For each co-assembly, genomic segments were clustered based on segment coverage vectors (mean and variance) using MetaBAT2[71] (version 2:v2.16-4-g40efa2d) with parameters “-s 1500 -m 1500 --maxP 95 --minS 60 --maxEdges 200 --seed 1 -l –saveCls”. The output was treated as initial genomic bins and trimmed as follows: Coverage vectors were computed separately for the two sides of each segment, over the interval starting 10bp away from segment border and up to 2000bp into the segment (or less for segments shorter than 2000bp); segment sides associated with a genomic bin were organized in a graph, where two sides were connected by an edge if they were associated with the same segment or if a comparison of their coverage vectors failed to separate them; and each connected component in the graph was then converted to a metagenome-assembled genome (MAG). In this manner some initial bins were split into several final MAGs. Each genome was associated with a unique population of a species in a specific subject.

**Dynamic variant classification.** The following procedure was applied to all true variants that were at least 200bp away from any segment boundary. To test if a variant $v$ is dynamic we compared 4 coverage vectors. We used the variant coverage vector $r_{var} = r(v)$, the local coverage vector $r_{local} = r(p)$, and the complement vector $r_{comp} = r_{local} - r_{var}$. Additionally, we defined the regional coverage vector $r_{region} = r(p - C, p + C)$, where $C$ equaled 1000bp or less if near an edge of the containing contig. If $r_{region}$ and $r_{local}$ were separated (P<0.01) we rejected the variant, since we require the regional and local coverage to be linked. We verified that $r_{var}$ and $r_{local}$ were separated (P<0.01); otherwise, we rejected the variant as non-dynamic, since it is either a result of paralogs within a genome or a polymorphic variant with a negligible contribution to fitness. We verified that $r_{region}$ and $r_{comp}$ were separated (P<0.01); otherwise, we rejected the variant as a possible result of ortholog sequences (sometimes referred to in the literature as recruited reads). Variants that passed all three tests were classified as dynamic variants. A variant was associated with a MAG if it was contained in one of the segments of the MAG.

### <u>Benchmarking the approach</u>

**Simulated communities.** We generated 100 random communities as follows. Let a genome be one or more sequences of nucleotides. Let $G$ be a set of genomes representing a community and let $n_{mut}$ be the number of designated mutations. To generate a random community, each genome $g \in G$ was associated with a population composed of two strains, where one strain $s_{g,1}$ was $g$ (called the baseline strain) and the second strain $s_{g,2}$ (called the mutated strain) was constructed by introducing $n_{mut}$ mutations to $g$ as follows. At each step, a random mutation was selected with a probability of 0.8 to be one of the 3 local mutations (substitution, insertion, or deletion) or otherwise to be one of the 3 global mutations (inversion, large insertion, or large deletion). A substitution was defined by a random position and a substituting nucleotide. A local insertion was defined by a random position in which a random nucleotide sequence (1-12nt long) was inserted.

A local deletion was defined by a random position at which 1-12nt were deleted. An inversion was defined by two positions that were fixed to be 1000bp apart and involved reversing the orientation of the sequence between the positions (i.e., transforming through a reverse-complement function). A large deletion was defined by a random 1000bp interval that was deleted. A large insertion involved the insertion of a mobile element randomly selected out of a pool of 4 randomly generated 1000bp mobile elements (same pool was used by entire community). To keep track of ground-truth through this process every mutation was logged alongside the 24bp identifier sequence that was upstream of the mutation (keeping a gap of 4bp), and the spanning interval of the mutation and the identifier were marked in a bitmask. When selecting a random position for a new mutation the bitmask representing previous mutations was used to make sure the new mutation that does not alter previous mutations or their identifiers. To model sequence-specific sequencing biases, non-overlapping genomic windows of 100bp were assigned random skew factors uniformly distributed between 1 and 2. The skew factor $f_{seq}(g, p)$ at position $p$ was determined through interpolation in the baseline genome. Factors were propagated in the mutated strain during the mutation process such that the baseline and mutated strains had matching factor profiles in syntenic regions. Strain genomes were then circularized, resulting in one circular chromosome per strain.

**Community datasets.** We generated 40 complex communities, each sampled with 30 genomes that were randomly selected from the set of Bacteroides and Firmicutes genomes in proGenomes2[72]. To these communities we applied one of 4 different mutation rates, $n_{mut} = 1, 10, 100, 1000$ (10 communities per mutation rate, 40 communities in total). We also generated 60 communities to examine the effect of genome relatedness by including in each community genomes from either a broad or a narrow taxonomic rank. We did so for 3 different cases: *Bacteroidetes* vs. the *Bacteroides* genus, *Firmicutes* vs. the *Clostridium* genus, and *Proteobacteria* vs. the *Enterobacteriaceae* family. For all cases, we generated 10 communities where 30 genomes were randomly selected from the broad rank and 10 communities in which genomes were limited to the narrow rank, while keeping the mutation rate fixed at 100 per genome (20 communities per case, 60 communities in total).

**Simulated abundance trajectories and sequencing factors.** Sixteen longitudinal samples were simulated for each community. For a community composed of genomes $G$, each genome $g \in G$ was assigned an abundance weight $\mu_g$, such that $\log_{10}(\mu_g)$ was uniformly distributed between 0 and 3. Sample-specific abundance weights $\mu_{g,i}$ for $i = 1, \dots, 16$ were normally distributed $\mu_{g,i} \sim \mathcal{N}(\mu_g, \mu_g)$ and restricted to the interval $[1,1000]$. The abundance $A_{g,i}$ of genome $g$ in library $i$ was set to $A_{g,i} = w_{g,i} / \sum_h w_{h,i}$. The abundance $A_{g,i,1}$ of the baseline strain $s_{g,1}$ in sample $i$ was set to $A_{g,i}$ for $1 \le i \le 8$ and otherwise set to $0.2 \cdot A_{g,i}$. Similarly, the abundance $A_{g,i,2}$ of the mutated strain $s_{g,2}$ in sample $i$ was set to 0 for $1 \le i \le 8$ and otherwise set to $0.8 \times A_{g,i}$.

**Sequencing bias.** Each strain $s$ was assigned a replication ratio $u_s$ that was uniformly distributed between 1 and 1.2, and the factor of library $i$ was normally distributed $u_{s,i} \sim \mathcal{N}(u_s, 0.33)$. Each position $p$ was assigned a replication bias factor $f_{rep}(s,p)$ based on a sinusoid that had a peak-to-trough ratio of $u_{s,i}$. Each position $p$ was assigned a final bias factor equal to $f(s,p) = f_{seq}(s,p) \times f_{rep}(s,p)$. Read probability at position $p$ was set to $P(s,p) = f(s,p) / \sum_q f(s,q)$.

**Simulated shotgun libraries.** Random paired reads (2x150nt) were generated for a community as follows. The total number of reads $R_i$ for the library of sample $i$ was set such that mean x-coverage across all genomes was 10x, taking into account differences in genome length. Each strain $s_{g,j}$ was assigned $A_{g,i,j} \times R_i$ reads in library $i$. Each read was assigned a position $f(s,p)$ by selecting a random position with probability $P(s,p)$. Sequenced molecule lengths were normally distributed $\mathcal{N}(400,10)$, enforcing a minimal length of 200. Strand was assigned randomly, and read pairs were generated from the strain genomes.

**Running simulated data.** The shotgun data of each simulated community were processed as described above for the real data, while skipping over the steps described in the processing raw reads section (adapter trimming, read quality filtering, removal of human reads). Briefly, reads were pooled to construct a community co-assembly, mapped back to the co-assembly, PolyPanner was applied, and the output was a set of MAGs and associated dynamic sites. For clarity, we distinguish between strain genomes (which were simulated) and MAGs (which are the output of PolyPanner). For each strain genome in the community, overlapping sequence intervals that were 100bp long (sliding windows with 10bp steps) were mapped to the co-assembly using BWA-MEM. Low quality alignments (edit distance >20, score >30, or alignment length >50) were discarded. Alignments were traversed to generate a 1bp mapping from co-assembly contig coordinates to zero or more genome coordinates. The entire co-assembly was divided into maximal alignment intervals by consolidating adjacent coordinates that are compatible, where each alignment interval $s$ perfectly aligns to zero or more strain genomes $G(s)$. For example, an interval $s$ that is a result of the assembly of two syntenic regions in strains $s_{g,1}, s_{g,2}$ is expected to align to both of them, or formally: $G(s) = \{s_{g,1}, s_{g,2}\}$. We define the alignment of a set of intervals $S$ to a set of genomes $G$ to be $I(S,G) = \{s \in S: G(s) = G\}$, or in other words, $I(S,G) \subseteq S$ is the subset of $S$ that perfectly aligns to all the genomes in $G$.

**Detection of assembly breakpoint.** Alignment intervals longer than 100bp were traversed in order along contigs, and pairs of intervals that aligned to a different set of genomes were marked as true assembly breakpoints. A reported assembly breakpoint was classified as true if the coordinate at which the breakpoint was identified was marked as a true breakpoint.

**Genome completeness and contamination.** Each MAG $b$, composed of alignment intervals $S_b$, was associated with a set of strain genomes $G_b = argmax_G|I(S_b, G)|$, or in other words, $G_b$ is the set of strain genomes that have the longest alignment to $b$. Completeness $C(b)$ was defined as $\frac{|I(S_b,G_b)|}{|G_b|}$, where $|G_b|$ is the average length of the genomes in $|G_b|$. In other words, $C(b)$ is the fraction of the genomes in $G_b$ which aligned to the MAG $b$. Contamination $X(b)$ was defined as $\frac{|X(S_b)|}{|S_b|}$, where $X(S_b) \subseteq S_b$ is the set of intervals in $S$ that align to genomes outside the set $G_b$.

**Variant detection.** Each source genome and associated MAG were processed as follows. Let $M$ be the set of introduced mutations, defined by their type and sequence identifier. Let $O$ be the set of observed variants for this MAG (referred to as 'true variants' above) and let $O_{dyn} \subseteq O$ denote the set of dynamic variants reported by the algorithm. For each mutation, we searched for the mutation identifier in the contigs of the MAG, and when there was a unique exact match, the mutation was associated with an expected variant that was generated based on the identity of the mutation, and in the precise location based on the position and orientation of the identifier in the co-assembly. This process resulted in a set of expected variants $E$. An observed and an expected variant were matched if they were identical (e.g., both involved a substitution of A for G) and the distance between their coordinates was zero for substitutions, up to 2 for indels and up to 4 for rearrangements. Genuine variants $O_{genuine} \subseteq O$ were defined as observed variants that had a matching expected variant. Spurious variants $O_{spurious} \subseteq O$ were defined as observed variants that lacked a matching expected variant and were also at least 200bp away from any segment edge. False variants $O_{false} \subseteq O_{dyn}$ were dynamic variants that lacked a matching expected variant.

Detected mutations $M_{detected} \subseteq M$ were mutations that had an associated expected variant that matched a dynamic variant. The density of spurious variants was defined as $|O_{spurious}|/L$, where $L$ is the total length of the contigs in the MAG. The percent of false detections was defined as $|O_{false}|/|O|$, or in other words this was the percent of variants that were reported as dynamic without a matching mutation. The percent of correctly reported variants (our measure of sensitivity) was defined as $|M_{detected}|/|M|$, and was similarly defined separately for each mutation type.

### Genome and variant annotation

**Metagenome-assembled genomes and their annotation.** All MAGs that were >500kb were assessed using CheckM[73] (v1.2.2, reference generated on 16/1/2015), which was run with the lineage_wf workflow using default parameters. The selected list of 5665 MAGs examined in this study were MAGs that were >50% complete and <10% contaminated. MAGs were taxonomically annotated using GTDB-Tk[74] (v2.2.6, reference database version R207_v2), using the classify_wf workflow with default parameters. 73 MAGs (1.28%) were resolved by GTDB-Tk down to the genus level (without reaching a species-level resolution) and were assigned a species by adding an "sp." suffix to the genus, e.g., "*Collinsella sp.*". 30 MAGs (0.52%) for which GTDB-Tk did reach a genus-level resolution were left without a species.

**Inference of strains.** To infer strains, Strain Finder[48] was applied to all MAGs that had between 1 and 1000 dynamic variants. Since the input of Strain Finder is solely nucleotides and we have additional types of variants (such as indels and rearrangements) we applied an encoding-decoding scheme, where for each polymorphic site the 2-4 variants at the site were encoded using arbitrary nucleotides (>99% sites were bi-allelic, no site had over 4 alleles), and site-specific conversion tables were used to decode nucleotides back to variants after Strain Finder terminated. Strain Finder (v1.0) was run with parameters "-e 1e-4 --n_keep 3 --max_reps 10 --dtol 1 --ntol 3 –converge", separately testing 2-8 strains, and the number of strains was selected using the Akaike information criterion (AIC). Each output strain was defined by a single variant per polymorphic site and a temporal frequency trajectory, with the frequencies of all strains of a MAG summing to 1 at each time point.

**Strain phylogeny tree and linkage groups.** For each MAG, strains were placed on a maximum parsimony tree using the function pratchet in the phangorn R package[75] (v2.11.1). The length of each tree branch was set to the number of sites that were inferred to change their state along the branch. Each variant $v$ was associated with a single branch $b(v)$ on which $v$ changed states. In case there were multiple branches on which $v$ changed states, a single branch with the minimum branch length was chosen. The set of variants associated with a branch is called the linkage group (LG) of the branch $V(b)$.

**Genome abundance trajectories.** The abundance of genome $g$ in library $i$ was defined to be $A_i(g) = r_i(g)/\sum_{g\in G} r_i(g)$, where $r_i(g)$ is the total number of reads covering genome $g$, and $G$ is the set of all genomes. The abundance trajectory of $g$ was $A(g) = (r_1(g), \dots, r_m(g))$, and the normalized abundance trajectory was $N(g) = A(g)/T(g)$, where $T(g) = \sum_{i=1,\dots,m} A_i(g)$. To generate **Fig. 1A**, normalized abundance trajectories were clustered using k-means (k=100) and sorted along the y-axis based on hierarchical clustering.

**Genes.** For each subject, genes were predicted with Prodigal[76] (v2.6.3), using the parameters "-p meta -g 11". Genes were blasted against the Uniref100 database (downloaded July 2020) with DIAMOND[77] (v2.0.15.153), using the 'blastp' command, assigning genes to top hits. Genes across all subjects, alongside *Escherichia coli* genes (K-12 MG1655, assembly ASM584v2), were clustered with MMseqs2[78] (version bdd169b3e285299cab792e62d60eb1f4e4e434d2), using parameters "--min-seq-id 0.5 -c 0.8 --cov-mode 0 --cluster-mode 0". Genes representative of clusters were annotated using the eggNOG-mapper[79] (emapper-2.1.7-bfd73c0, reference database 5.0.2), using parameters "--itype proteins". We focused on the KEGG Orthology (KO) of genes, as reported by eggNOG. Note that some genes were annotated by eggNOG with multiple KOs. There were 936 gene clusters (representing 21544 genes) initially annotated as K02469 (*gyrA*) and/or K02621 (*parC*). These genes were reclassified as K02469 if their eggNOG name was '*gyrA*', the remaining genes were reclassified as K02621 if they matched the PFAM entry 'DNA_topoisoIV'; genes meeting neither criterion were dropped from downstream analysis. After the reclassification, there were 11777 genes annotated as *gyrA* with K02469 and 9727 annotated as *parC* with K02621. A gene was associated with a MAG if it was completely contained in one of the segments of the MAG. Genes not associated with any of the 5665 MAGs were dropped from downstream analysis. Variants were classified as intra-genic if contained within a gene and otherwise classified as inter-genic, and each was associated with the genes that were upstream and downstream of the variant, if present.

**Sweeping variants.** The average frequency of variant $v$ at position $p$ over samples $I$ was defined to be $\sum_{i \in I} r_i(v) / \sum_{i \in I} r_i(p)$, where $r_i(v)$ is the number of reads supporting the variant in sample $i$ and $r_i(p)$ is the number of reads supporting position $p$ (i.e., all variants) in sample $i$. Variants that had a frequency above 50% in the baseline samples (days -2 to 0) were reversed (e.g., "A to T" was transformed to "T to A"). A variant was classified as sweeping if it had an average frequency <20% in the baseline samples and an average frequency >80% in the post-antibiotic samples (days 10-28). To determine if a genome had sufficient coverage to detect sweeps, an artificial variant trajectory that sweeps from a frequency of 0% to 100% as of day 10 and with a total x-coverage based on the genome x-coverage trajectory was tested using the same statistical tests that were applied to all variants (namely the ortholog and paralog tests, defined above). All downstream analysis was limited to sweeping variants that were part of small LGs (up to 100 variants/LG).

### Analysis of evolutionary dynamics

**Parallel evolution analysis.** We assigned every LG a weight of 1 and equally distributed the weight between all genes associated with one or more variants in the linkage group. Gene weights were distributed between all gene KOs (weight dropped if no KO was associated). KO total weights were computed by summing over the LGs. A background weight distribution was generated by creating $10^6$ random sets of variants, by replacing the genes of an LG $V$ with a random set of $|V|$ genes uniformly selected from the genes of the MAG associated with $V$. The $p$-value of each KO was empirically calculated by embedding the observed weight in the distribution of random weights. KO enrichment ratios were computed by dividing the observed weight and the mean expected weight. We considered only KOs that had a $p$-value below 0.05, an enrichment ratio of at least 2-fold, and for which the associated supporting variants were found in at least 3 different subjects. False discovery rates ($q$-values) were added using the Benjamini-Hochberg approach.

***GyrA* analysis.** Genes annotated as *gyrA* (K02469), including the *E. coli* reference gene, were aligned with Clustal Omega[80] (v1.2.4), using default parameters. For each variant, the *E. coli* coordinate was set to the closest *E. coli* coordinate according to the global alignment of all genes.

There were 4987 MAGs that had a *gyrA* gene. The amino acid at position *gyrA*:83 (as shown in **Fig. 3C**) is shown for 698 MAGs that (1) had a single *gyrA* that aligned to the *E. coli gyrA* at position #83, and (2) had sufficient coverage to detect sweeps, if present (defined in section 'Sweeping variants' above). There were 56 MAGs in which *gyrA*:83 changed identity due to a sweeping substitution variant. Species-specific resistance alleles at position *gyrA*:83 were defined based on the substituting amino acids of the 56 substitutions at *gyrA*:83.

**Evolvability analysis.** For this analysis we focused on 410 populations that had serine at *gyrA*:83 and had sufficient coverage for detection of sweeps, if present (defined in section 'Sweeping variants' above). We trained models to predict two response variables: *gyrA evolvability*, defined as the probability of the population to undergo one or more sweeps involving gyrA, and non-*gyrA evolvability*, defined as the probability of the population to undergo one or more sweeps involving any gene except *gyrA*. As predictor variables we used the baseline abundance ('Base', days -2 to 0), the abundance during antibiotics ('Treated', days 1-5), the abundance post-antibiotics ('Post', days 10-28), and the abundance at last sample ('Late', day 77). All abundance values were log-transformed after adding 0.001%. Additional variables were also considered: the fold-decrease in abundance during antibiotics ('Decline', equal to $\log_{10}$(Base/Treatment)) and 2 phylum variables. Separately for the two response variables, we trained 9 logistic regression models (Base, Treated, Post, Late, Decline, Base+Decline, Base+Phylum, Decline+Phylum, Base+Decline+Phylum), with k-fold validation using the caret package in R, with the 'repeatedcv' method (k=10 and 10 repeats). We rejected models if one of the coefficients was not significant (using a threshold *p*-value of 0.05). Models were ranked based on their Akaike information criterion (AIC). The pROC package in R was used to plot ROC curves (receiver operating characteristic curves) and compute the area under the curve (AUC) for all models.

**Recovery analysis.** Analysis was performed on all 1771 sweeping variants. We inferred a selection coefficient separately for each sweeping variant under the simplistic assumption that selection coefficients are constant over time, and using a maximum likelihood approach as follows. The relative frequency of the variant over time equals $\frac{p(t)}{1-p(t)} = c \cdot (1-s)^t$ (equation 1), where $p(t)$ is the frequency of the sweeping variant at generation $t$, and $s > 0$ is the selection coefficient, representing the fitness advantage of the baseline variant state compared to the swept state [81]. The data are a sequence of triplets $D = (k_i, n_i, d_i)_{i=1}^{N}$, where $N = 4$ is the number of post-antibiotic samples (sampled on canonical days 10, 18, 28, 77), $k_i$ is the number of reads supporting the variant, $n_i$ is the number of reads supporting the variant position, and $d_i$ is the actual sampling day of sample $i$. Based on equation 1, we define $p_i = \frac{c \cdot (1-s)^{d_i \cdot m}}{1+c \cdot (1-s)^{d_i \cdot m}}$, where $m = 10$ is the number of generations per day. We model the probability of the observed data at sample $i$ using a binomial function: $P(X_i = k_i) = \binom{n_i}{k_i} {p_i}^{k_i}(1-p_i)^{n_i - k_i}$, where $X_i$ is a random variable representing the number of reads supporting the variant at sample $i$. The selection coefficient $s$ and the initial ratio $c$ were selected to maximize the likelihood function $L(s, c|D) = \prod_{i=1}^{N} P(X_i = k_i)$, using the L-BFGS-B method in the optim function in R, constraining $-0.4 < s < 0.4$ and $10^{-6} < c < 10^{6}$, and initializing $s_0 = 0$ and $c_0 = 1$. The optimization converged for 1470 variants (61 of which were associated with *gyrA*). Given optimized $s$ and $c$, we calculated the number of days until the frequency reached 1% using equation 1 above. Note that the number of generations per day $m$ scales the selection coefficients but does not affect the number of days until recovery.

### Supplementary Text

**Supplementary Note 1.** The number of resistant cells is estimated to equal $\mu \times N \times \frac{(1-e^{-ts})}{s}$, where $\mu$ is the mutation error rate (mutations per bp per generation), $N$ is the total number of cells (i.e., population size), $t$ is the number of generations since the last sweep or colonization event, and $s$ is the selection coefficient representing the fitness cost of the resistant allele while there is no antibiotic exposure (see Eq. 3.9 in ref.[82] that deals with the case in which $t = \infty$ and and Eq. 7 in ref.[50]). We assume $s \leq 0.01$ (in line with results in this work), $t \geq 70$ (indicating at least one week passed since the last selective sweep), and that there are $10^{13}$ cells in the intestine of a subject. Requiring at least one resistant cell results in an upper bound on the population abundance threshold that equals 0.0002% and 0.002%, for $\mu = 10^{-9}$ and $\mu = 10^{-10}$ respectively.

### Supplementary Table Legends

**Supp. Table S1. Genome table.** Information on the 5665 genomes described in this study. Table columns:

gid: genome identifier.

aid: subject identifier.

bin: internal genome identifier.

xcoverage: mean x-coverage of genome.

length: genome length (bp).

n_strains: number of strains.

complete: genome completeness.

contam: genome contamination.

is.detected: does genome have enough x-coverage to detect sweeps.

strain.class: strain classification.

K02469_83_value: baseline value at *gyrA*:83.

K02469_83_mut: substitution at *gyrA*:83.

vars: number of dynamic variants.

var.genes: number of unique genes associated with dynamic variants.

sweep.vars: number of sweeping dynamic variants.

sweep.genes: number of unique genes associated with sweeping dynamic variants.

phylum/class/order/family/genus/species: taxonomic identity.

**Supp. Table S2. Dynamic variant table.** Description of dynamic variants. For intra-genic variants gene_1 is the containing gene. For inter-genic variants gene_1 and gene_2 are the two adjacent genes. Table columns:

gid: genome identifier.

xid: variant identifier.

aid: subject identifier.

contig/coord: variant position.

variant: variant description.

edge_size: size of associated linkage group.

response: is variant sweeping.

K02469_83: coordinate within gene of *gyrA*:83, if gene is *gyrA*.

gene_1/2: gene identifier.

orient_1/2: orientation relative to gene.

uniref_1/2: Uniref100 identifier.

identity_1/2: Uniref100 sequence identity.

prot_desc_1/2: Uniref100 protein description.

start_dist_1/2: distance of variant from gene transcription start site (TSS).

mut_class_1/2: type of mutation.

mut_label_1/2: mutation label.

KEGG_ko_1/2: gene KO.

PFAMs_1/2: gene PFAM.

**Supp. Table S3. Genomes used to annotate *gyrA* and *parC*.** Table with 120 annotated reference genes that were used to validate the annotation approach of *gyrA* and *parC*. Table columns:

Index: running index.

accession: NCBI accession identifier.

desc: gene description.

taxa: taxonomic identity.

class: gene class, based on description and paper describing gene, if present.

**Supp. Table S4.** Table of KOs that showed evidence of convergent evolution. Table columns:

feature: KO identifier.

description: KO description.

pvalue: *p*-value of KO.

qvalue: *q*-value of KO computed using the Benjamini Hochberg correction.

enrichment: weight enrichment ratio of observed weight over an expected weight derived through permutations.

weight: observed total weight.

variant.count: number of variants associated with KO.

vc.count: number of unique linkage groups associated with KO.

bin.count: number of unique genomes associated with KO.

assemblies.count: number of unique subjects associated with KO.

median.vc.size: median linkage group size of associated variants.

genic.fraction: fraction of genic variants associated with KO.

Ns/Nn/Ks/Kn: statistics used to compute dN/dS ratios.

dNDs: dN/dS ratio of KO.
