## Supplementary material for "A short course of antibiotics selects for persistent resistance in the human gut": Supp. Figures

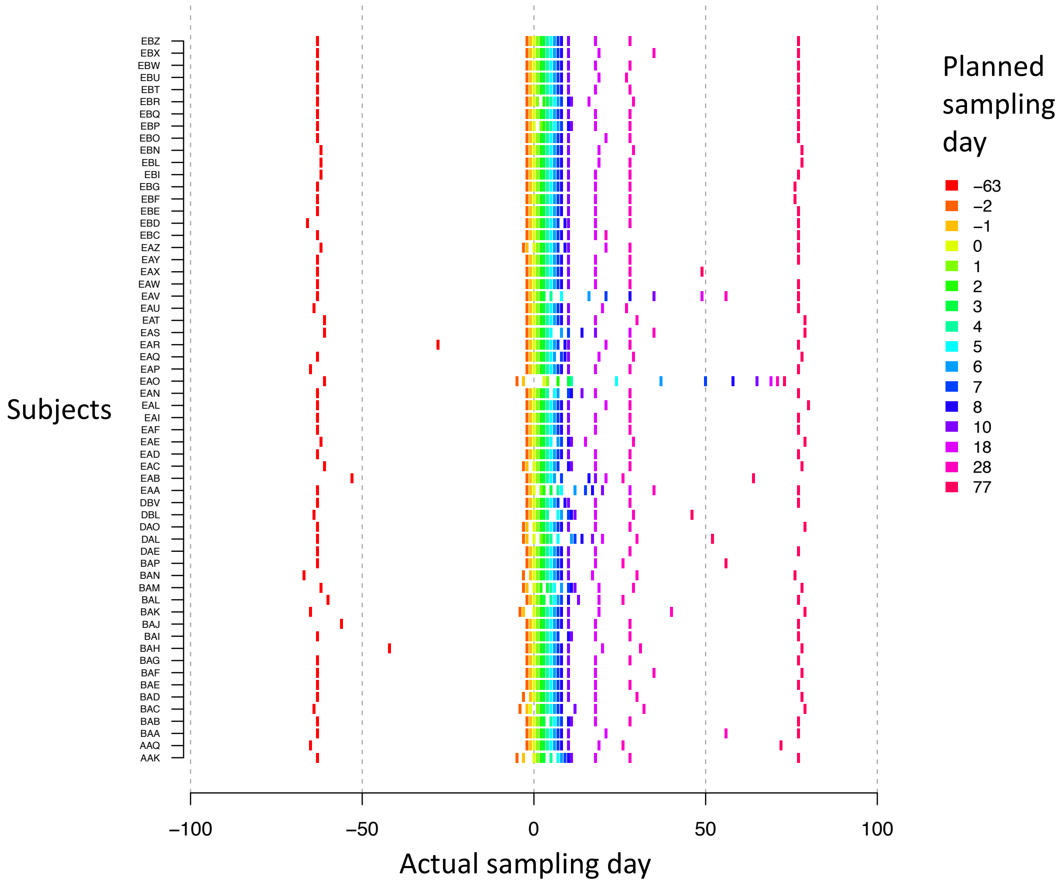


**Fig. S1. Sampling days.** The planned and actual sampling day for the 960 samples presented in this study. Shown is the actual sampling day (x-axis, relative to start of antibiotics) and subject (y-axis), colored by planned sampling day. Antibiotics were taken on days 0-4. When a planned sample was not available (not collected by subject or had technical issues in the extraction and sequencing process) all subject samples were shifted, while minimizing temporal changes. 76% of the actual samples were exactly as planned and 94.6% were up to 3 days away from the planned sampling day.

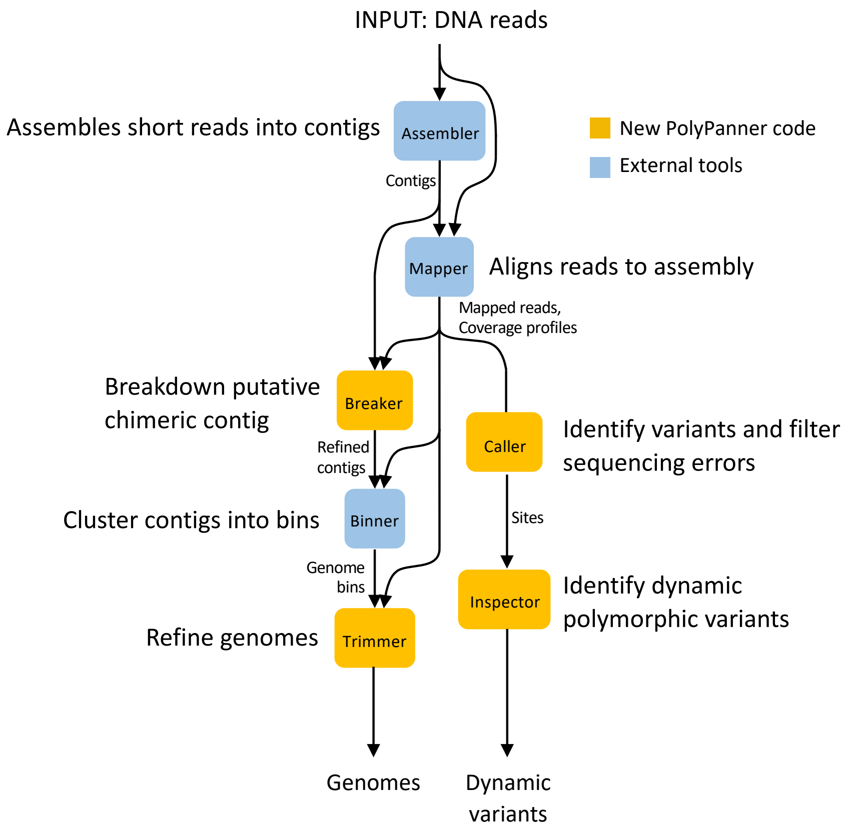


**Fig. S2. Overview of the PolyPanner workflow.** Each community (i.e., subject) is processed independently. Assembler) Sequenced DNA libraries from temporal samples are pooled and assembled. Mapper) Each DNA library is mapped to the assembly, generating 1-nt coverage profiles. Breaker) Contigs are refined into segments with consistent coverage profiles. Binner) Refined contigs are binned into genome bins. Trimmer) Genome bins are trimmed based on coverage profiles. Caller) Alignment mismatches of reads are counted for each genomic position, sequencing errors are modeled and removed, and variants are called in the remaining segregating positions. Inspector) Variants are classified according to coverage profiles, outputting dynamic variants. Final output is a set of genomes, each associated with set of dynamic variants, i.e., genuine polymorphic sites.

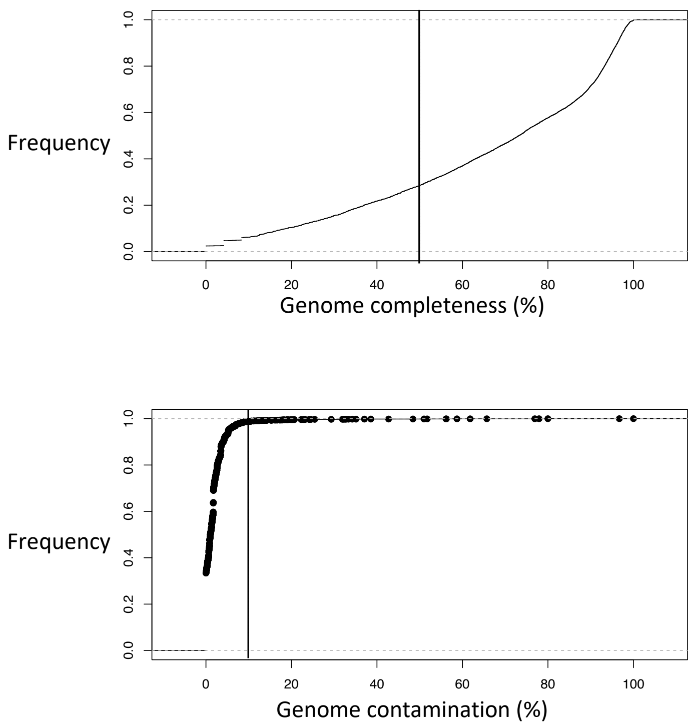


**Fig. S3. Genome quality.** Genome quality was assessed according to single-copy genes. An eCDF (empirical cumulative distribution function) is shown for genome completeness (top) and genome contamination (bottom). The completeness threshold (50%) and contamination threshold (10%) used to select the 5665 genomes presented in this study are depicted with black vertical lines.

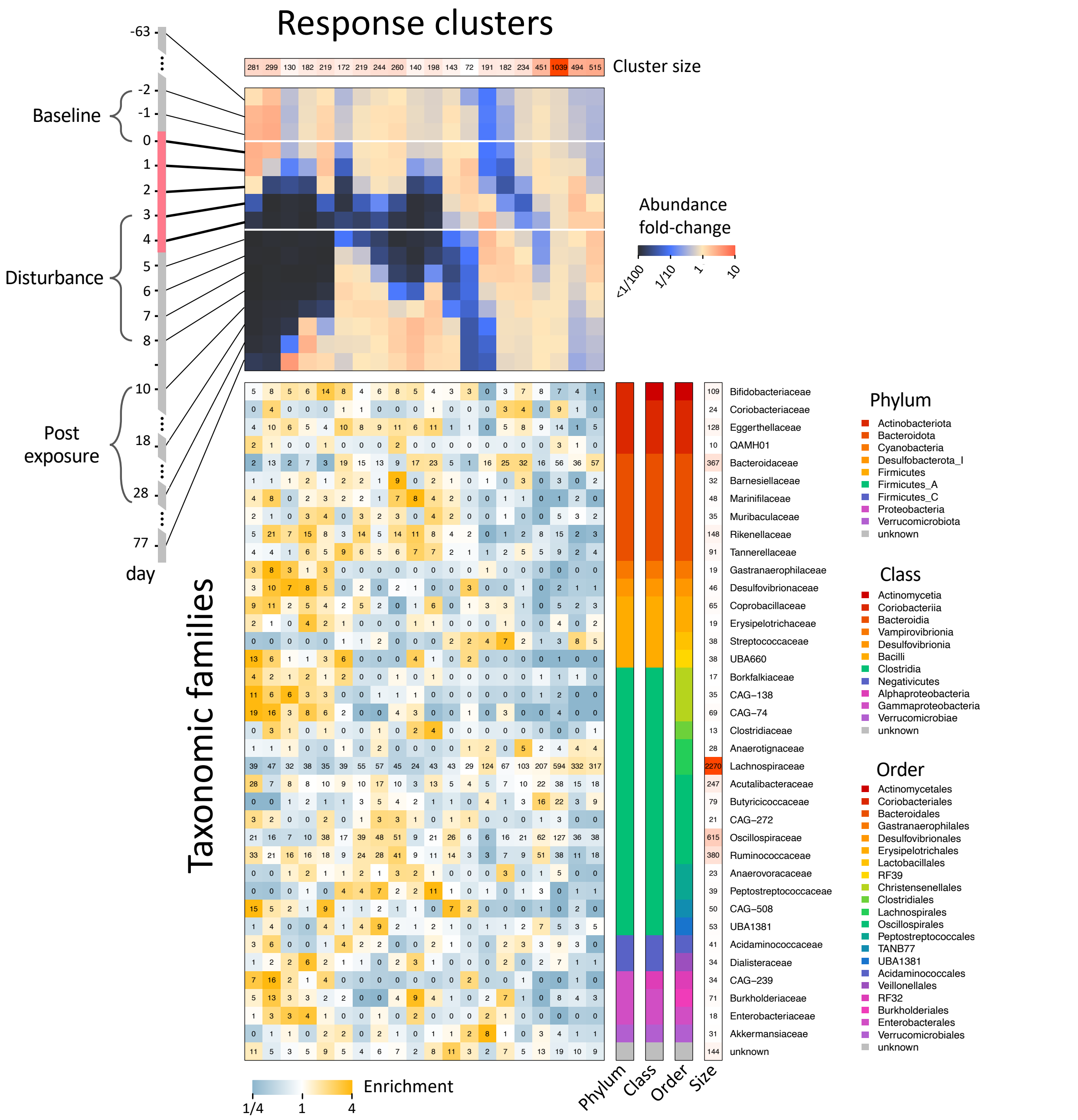


**Fig. S4. Association between response trends and microbial taxonomy.** Populations were clustered into 20 response trends (shown on top) and were associated with genera using GTDB-tk (shown on right). Only families associated with at least 10 populations are shown. The matrix (middle) is colored according to the enrichment of the number of populations, relative to an expected value assuming taxonomy and response trends are independent. Numbers in squares indicate the number of populations. Marginal number of populations colored in shades of red. Families differ in their response trends. For example, most *Akkermansiaceae* populations are blooming while most *Enterobacteriaceae* populations are dropping during the disturbance.

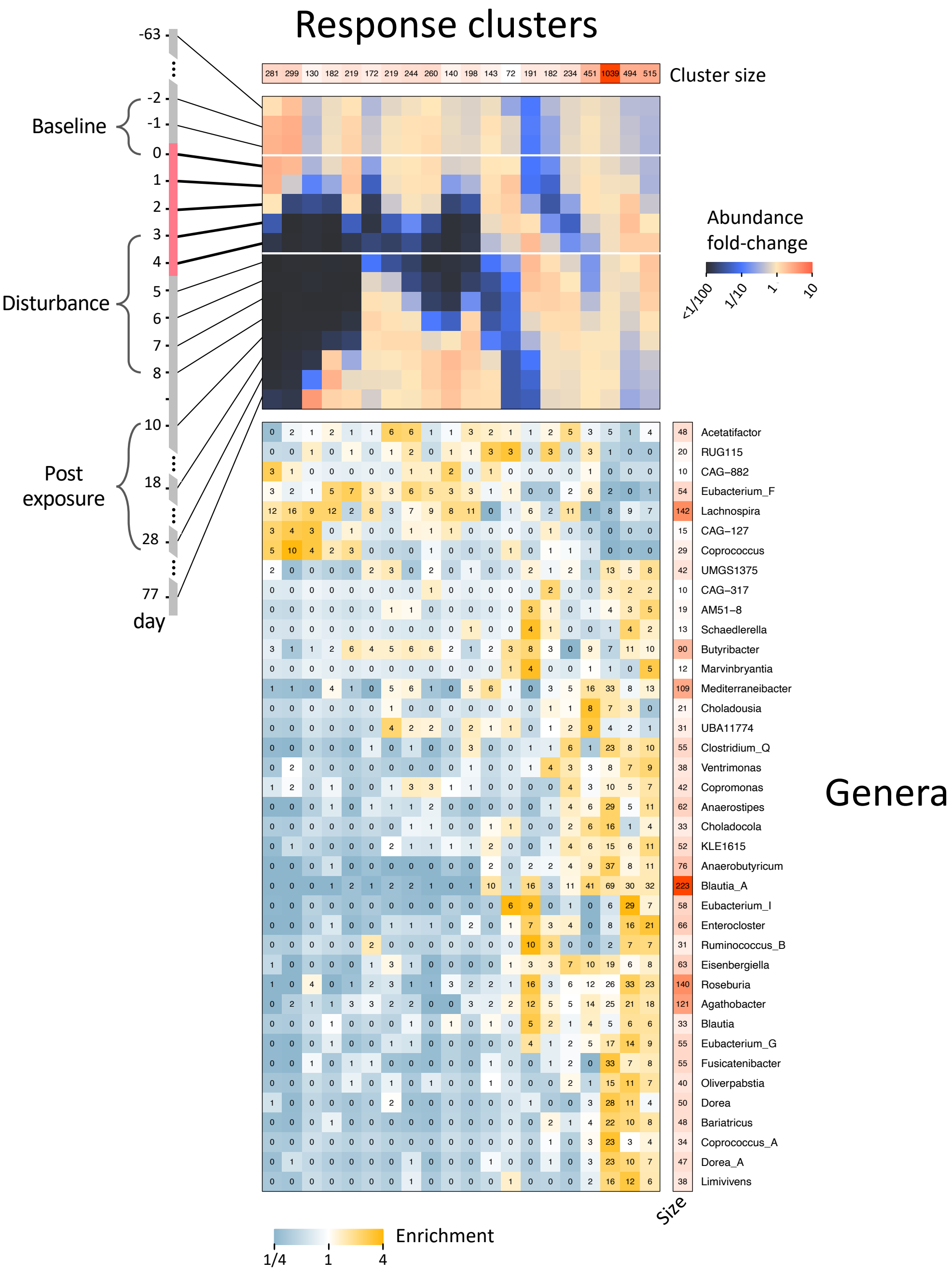


**Fig. S5. Diverse response trends within the *Lachnospiraceae* family.** Same as Figure S4, focusing on genera within the *Lachnospiraceae* family with at least 10 populations. Genera sorted according to hierarchical clustering of genera response enrichment vectors. Genera differ in their response to the disturbance. Note for example, how *Lachnospira* collapses and *Eubacterium_I* blooms.

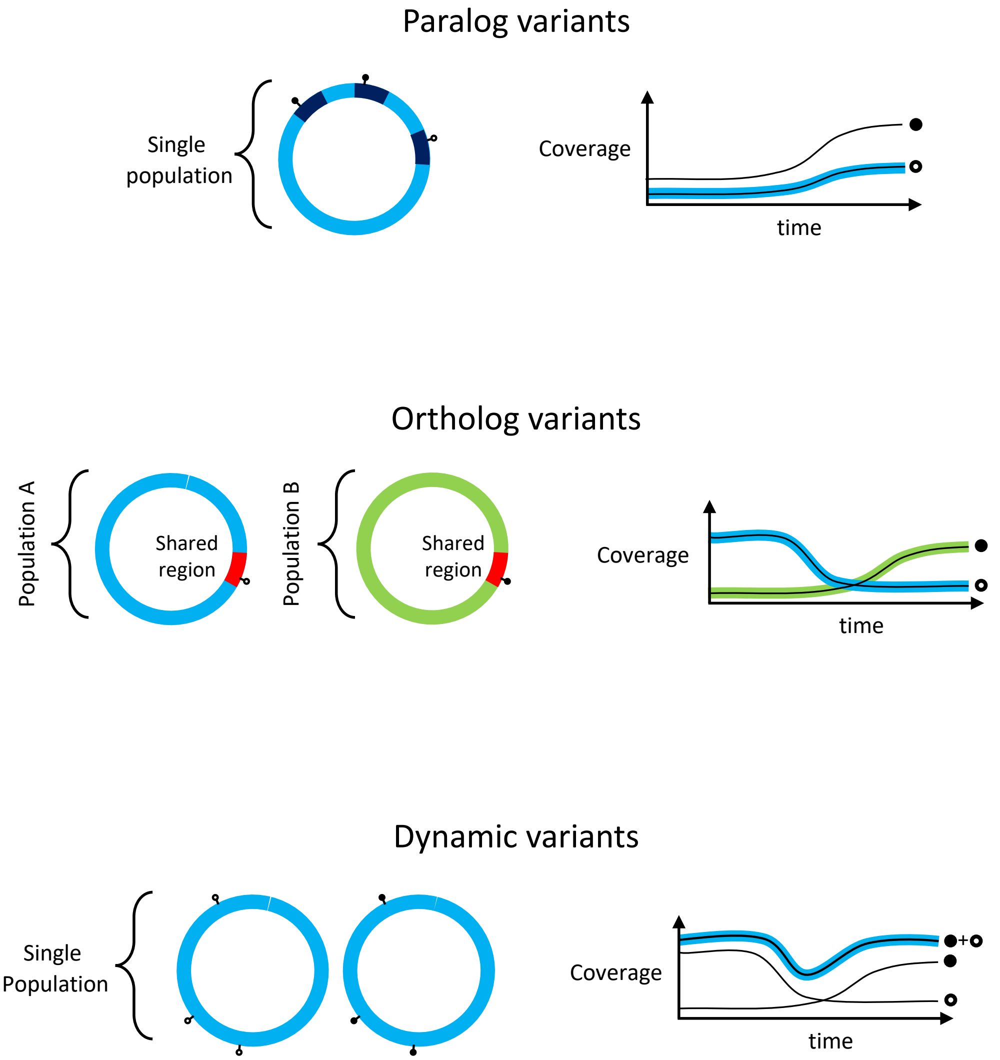


**Fig. S6. Classification of polymorphic variants.** We compute for each variant site three temporal coverage vectors: $\boldsymbol{M}$ (major allele, filled circles), $\boldsymbol{m}$ (minor allele, empty circles) and $\boldsymbol{R}$ (regional coverage, a proxy of the population coverage, colored lines). Each variant is classified in the following order. A variant is classified as a *paralog* if $\boldsymbol{M}$ and $\boldsymbol{m}$ are dependent (top). It is classified as an *ortholog* if $\boldsymbol{M}$ and $\boldsymbol{R}$ or $\boldsymbol{m}$ and $\boldsymbol{R}$ are dependent (middle). It is classified as *dynamic* if $\boldsymbol{M} + \boldsymbol{m}$ and $\boldsymbol{R}$ are dependent (bottom). For each scenario we show a possible genome configuration (left) and the coverage vectors over time (right). Downstream analysis in this work is limited to dynamic variants. Dependency is assessed using a chi-square test, see methods for complete details.

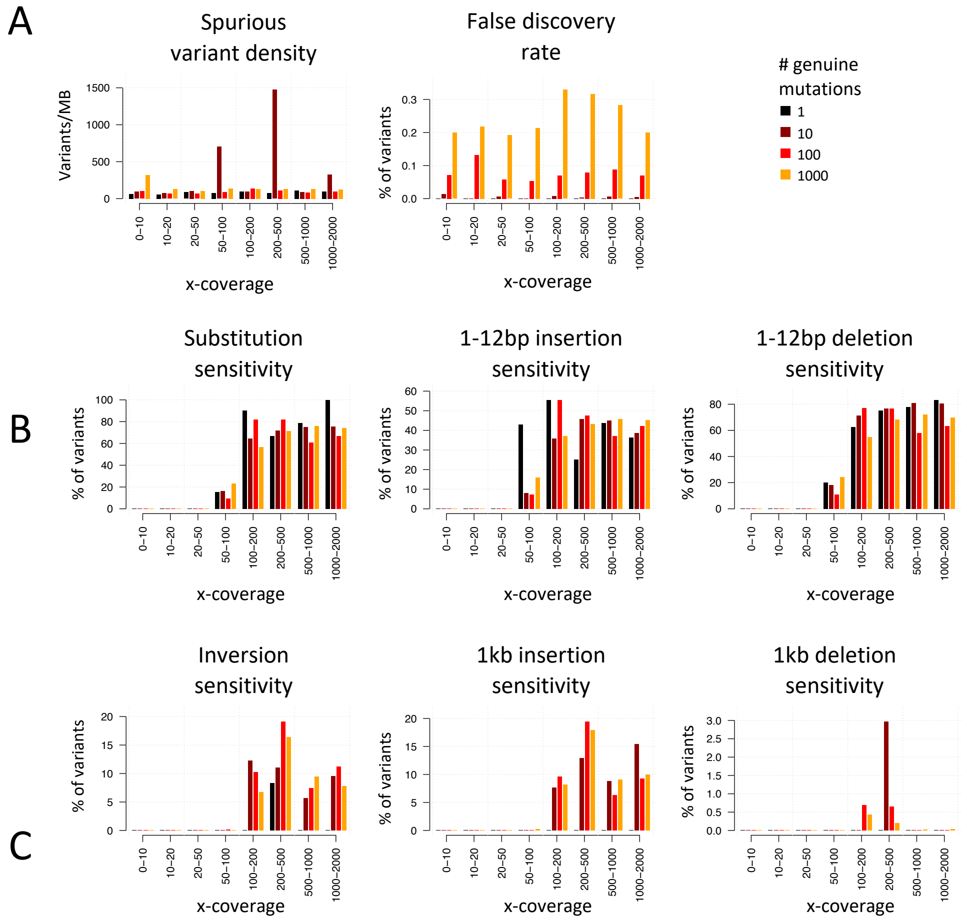


**Fig. S7. Performance on simulated communities.** Temporal sequencing data was simulated for 40 communities, divided into 4 groups according to the density of introduced mutations (1,10,100 or 1000 per genome), with 10 random communities per group. For each community, 30 genomes from the *Bacteroidetes* and *Firmicutes* phylum were randomly selected from the proGenomes database. For each genome, mutations were introduced into a simulated minor strain resulting in a ground truth of known genuine polymorphic variants. The mutation type was randomly selected among single nucleotide substitutions, short insertions and deletions (1-12bp), inversions with breakpoints that were 1000bp apart, and long insertions and deletions that involved 1000bp of DNA. Each community had 16 temporal samples, with the minor strain absent in samples 1-8 and at a frequency of 80% in samples 9-16. Genomes had a random x-coverage ranging 0-2000 reads per bp. In all panels, genomes are stratified by their x-coverage (x-axis) and the number of introduced mutations (colors). A) From left to right: Density of spurious variants (i.e., variants not associated with introduced mutations), percent of variants misclassified as dynamic out of all variants. B) Percent of introduced mutations that were correctly detected for substitutions, short insertions and deletions. C) Percent of introduced mutations that were correctly detected for inversion breakpoints, long insertions and long deletions.

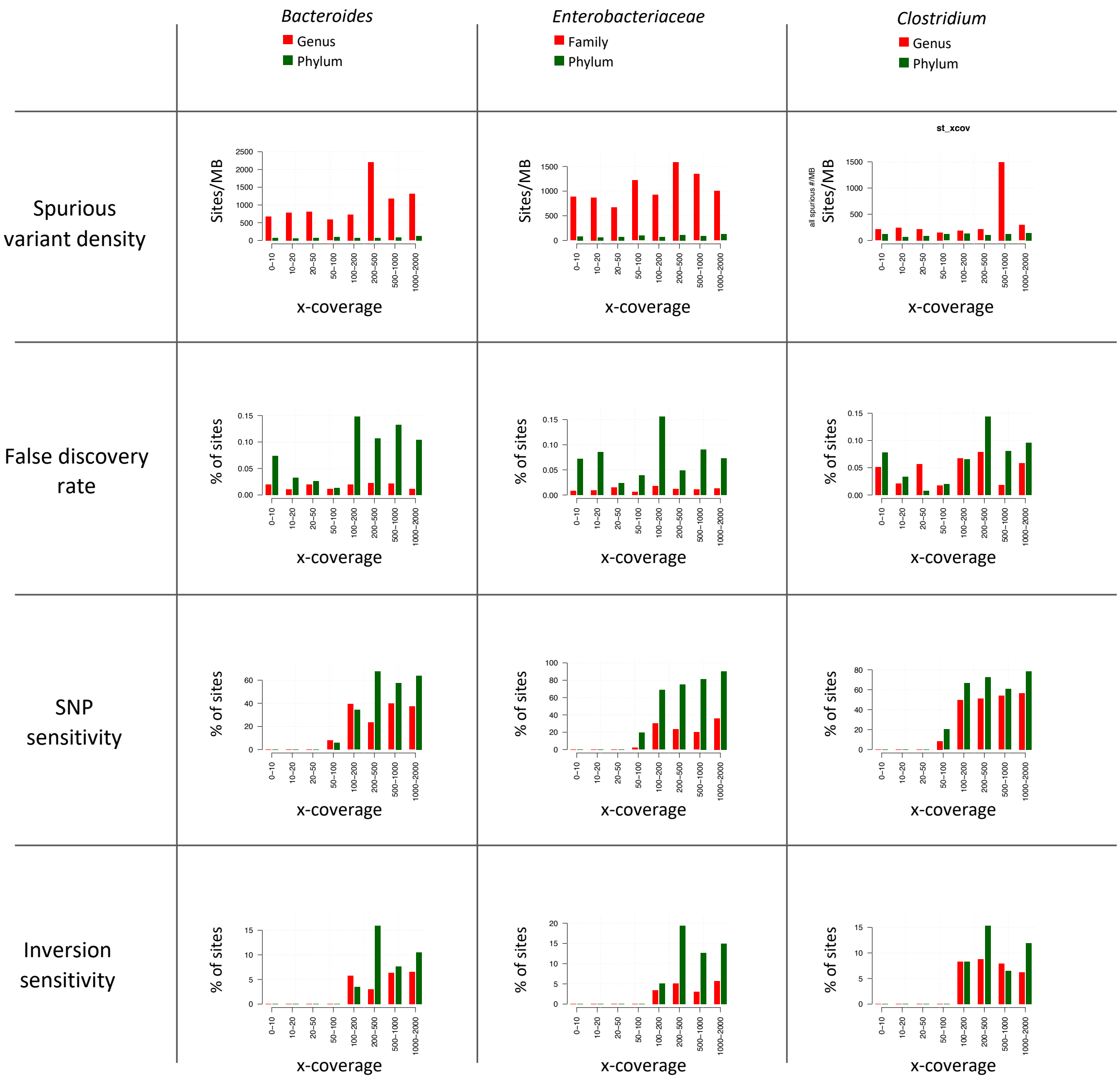


**Fig. S8. Genome crowding.** The presence of closely related genomes exacerbates the density of spurious variants and lowers sensitivity for detecting spurious variants in a taxonomy-dependent manner. We compared 10 communities composed of genomes limited to a genus or family vs. 10 communities sampled from the entire phylum for 3 cases: *Bacteroides* genus vs. the entire *Bacteroidetes* phylum (left), the *Enterobacteriaceae* family vs. the *Proteobacteria* phylum (middle), and the *Clostridium* genus vs. the *Firmicutes* phylum (right). Generation of minor strains and simulated reads as for Supp. Fig. 7, while introducing 100 mutations per genome. The density of spurious variants was elevated with genetic crowding for *Bacteroides* and *Enterobacteriaceae*, but not for *Clostridium*, a diverse genus. False discovery rate was below 0.15% for cases. Sensitivity varied between clades.

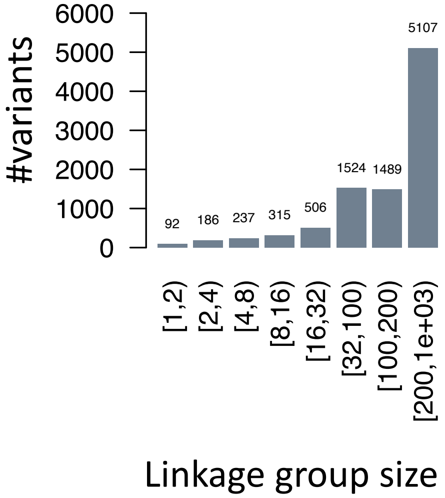


**Fig. S9. Linkage group size.** Histogram shows the distribution of sweeping dynamic variants by the size of their linkage group.

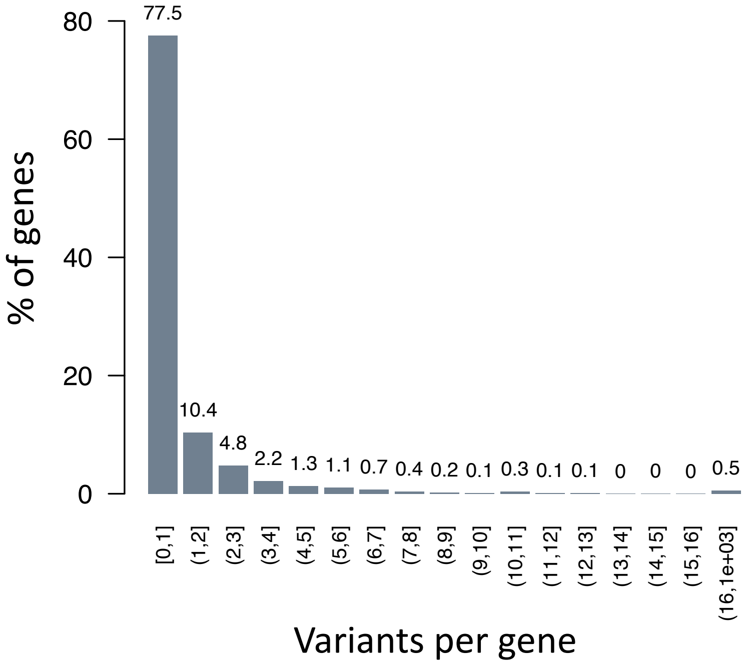


**Fig. S10. Variants per gene.** Histogram shows the percentage of 2016 genes that were in or adjacent to 1771 sweeping variants, according to the number of sweeping variants that were in or adjacent to them. These sweeping variants are associated with small linkage groups of no more than 100 variants, so the average density of variants across the genome is low. Nonetheless, 22.5% of genes associated with any variant were associated with more than one variant, indicating that horizontal gene transfer and recombination may have introduced multiple variants to these genes.

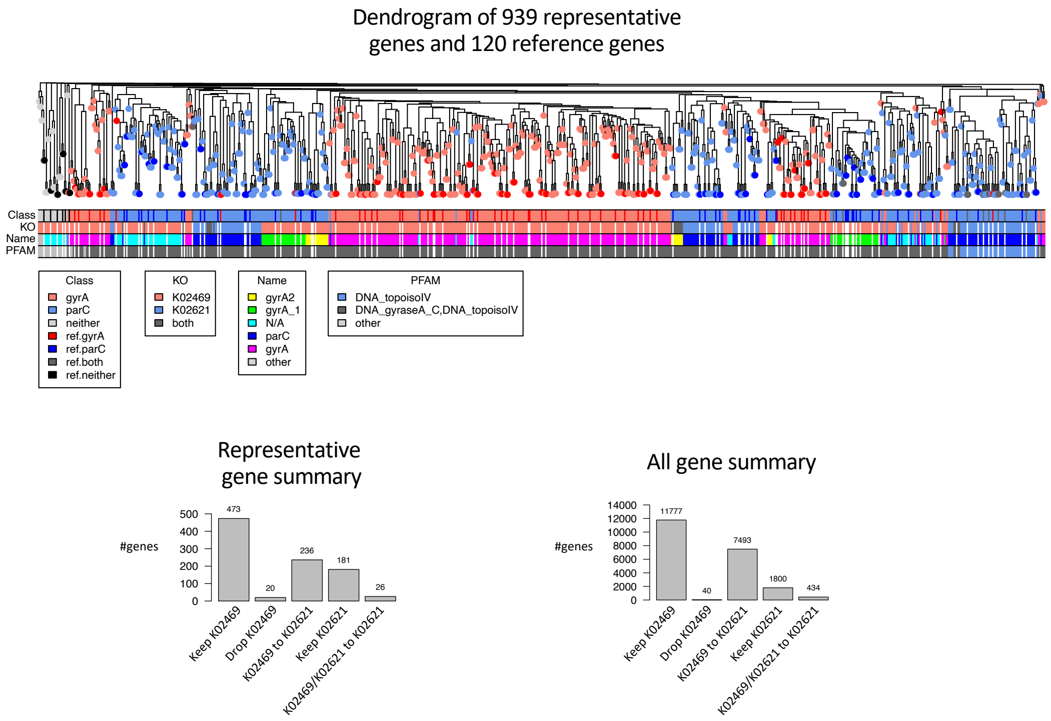


**Fig. S11: Separating the homologs *gyrA* and *parC*.** Genes were clustered using mmseqs2, and cluster representatives were annotated using eggNOG. There were 936 topoisomerase IIa associated gene clusters (representing 21544 genes) initially annotated as K02469 (*gyrA*) and/or K02621 (*parC*). Genes were reclassified as K02469 if their eggNOG name was '*gyrA*', the remaining genes were reclassified as K02621 if matching the PFAM entry 'DNA_topoisoIV', and genes meeting neither criterion were dropped. Topoisomerase IIa genes were analyzed alongside 120 annotated reference genes downloaded from NCBI (Supplementary Table S3) to validate the annotation approach. Shown is a dendrogram of genes >100aa clustered based on amino acid sequence alignment identity and colored according to gene classification (reference genes shown with darker colors). Shown below, from top to bottom, are final classifications, original KO assignments, eggNOG names and the PFAM hits. There is a near-perfect correspondence between the new annotations and annotations of reference genes, validating our reclassification approach. Barplots shown below summarize the number of genes that kept or switched annotations (left: representative genes, right: all genes). After reclassification, there were 11777 genes annotated as *gyrA* with K02469 and 9727 annotated as *parC* with K02621.

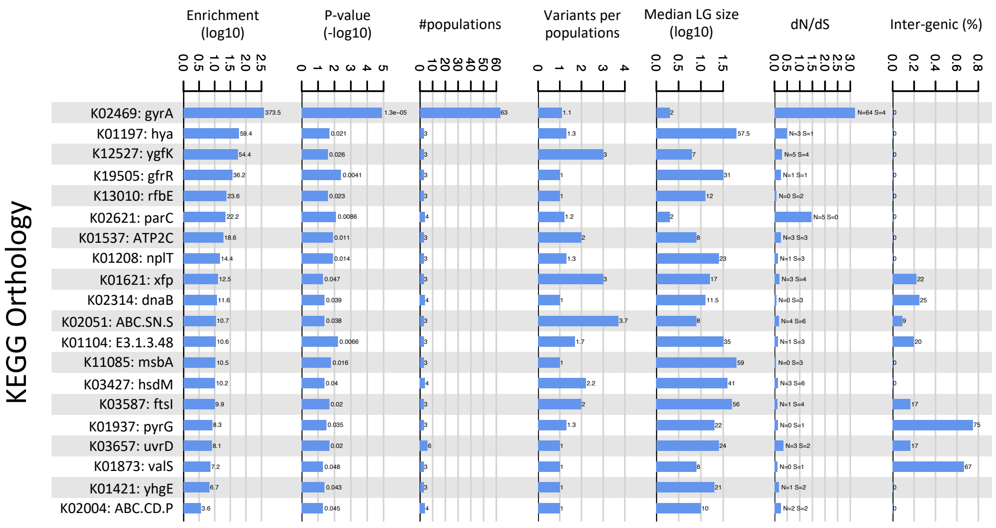


**Fig. S12. KEGG orthology table.** All 20 KEGG orthology entries (KOs) significantly enriched in sweeping variants are shown. Columns describe attributes of the variants associated with the KO, from left to right: Observed over expected enrichment ratios (background model generated through shuffling of variants within their respective genomes. Enrichment *p*-values. Number of populations with one or more KO-associated variants. Average number of KO variants per population. Median size of linkage group. dN/dS ratios of assigned variants, calculated from intragenic substitution variants. Percent of intergenic variants. For visualization purposes dN/dS ratios were calculated using a pseudo count of 1: $dN/dS=[(N+1)/(S+1)]/[(n+1)/(s+1)]$, where N and S are the observed number non-synonymous and synonymous, and n and s are the possible number non-synonymous and synonymous, respectively.

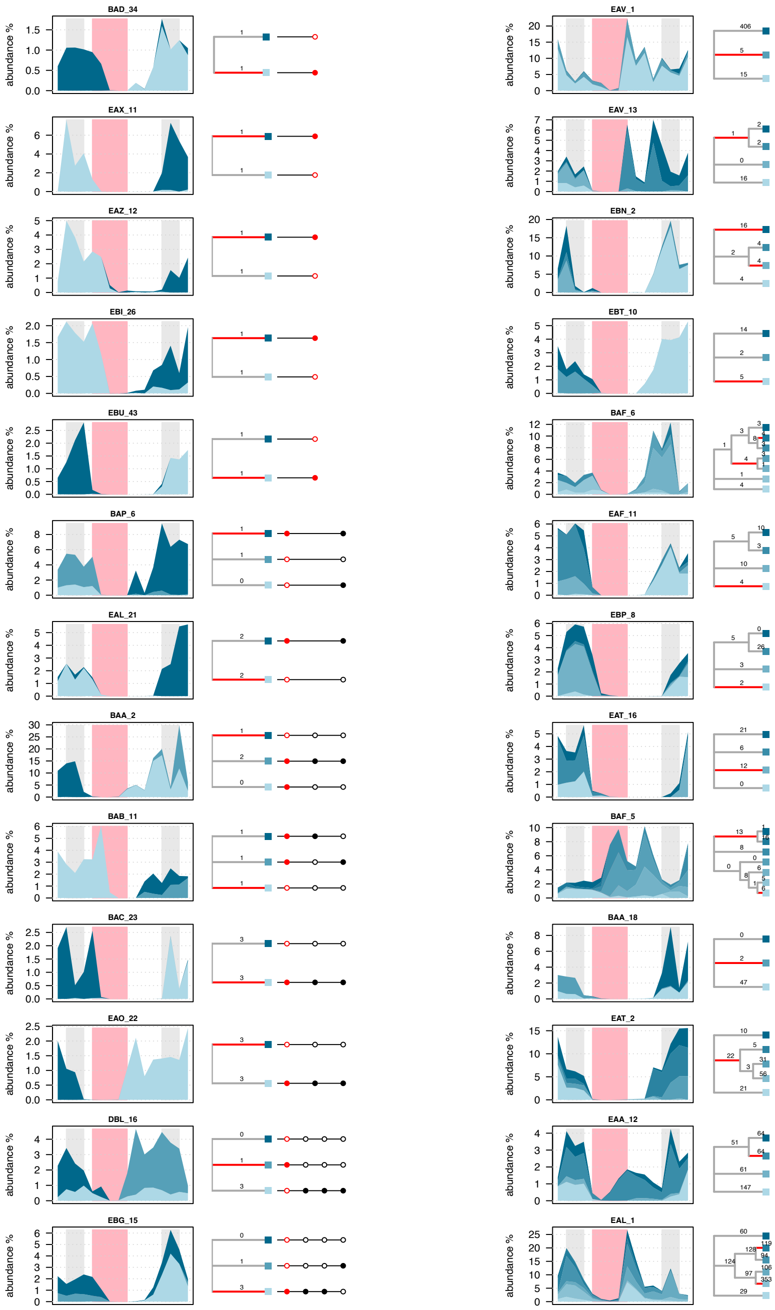


**Fig. S13. Examples of evolving populations associated with a mutation in *gyrA*.** For each population shown from left to right are the the strain abundance plot, the strain phylogeny tree (number above each branch depicts the size of the associated linkage group, colored red if branch associated with a change in *gyrA*), and the variant-strain matrix (*gyrA* variants colored red). In the strain abundance plots, abundances of strains are stacked, background colors emphasizing baseline samples (days -2 to 0), disturbed samples (days 3-8) and post-exposure samples (days 10-28). Strains are colored in shades of slate gray and organized top to bottom in a consistent order. See Supp. Table S2 for variant details.

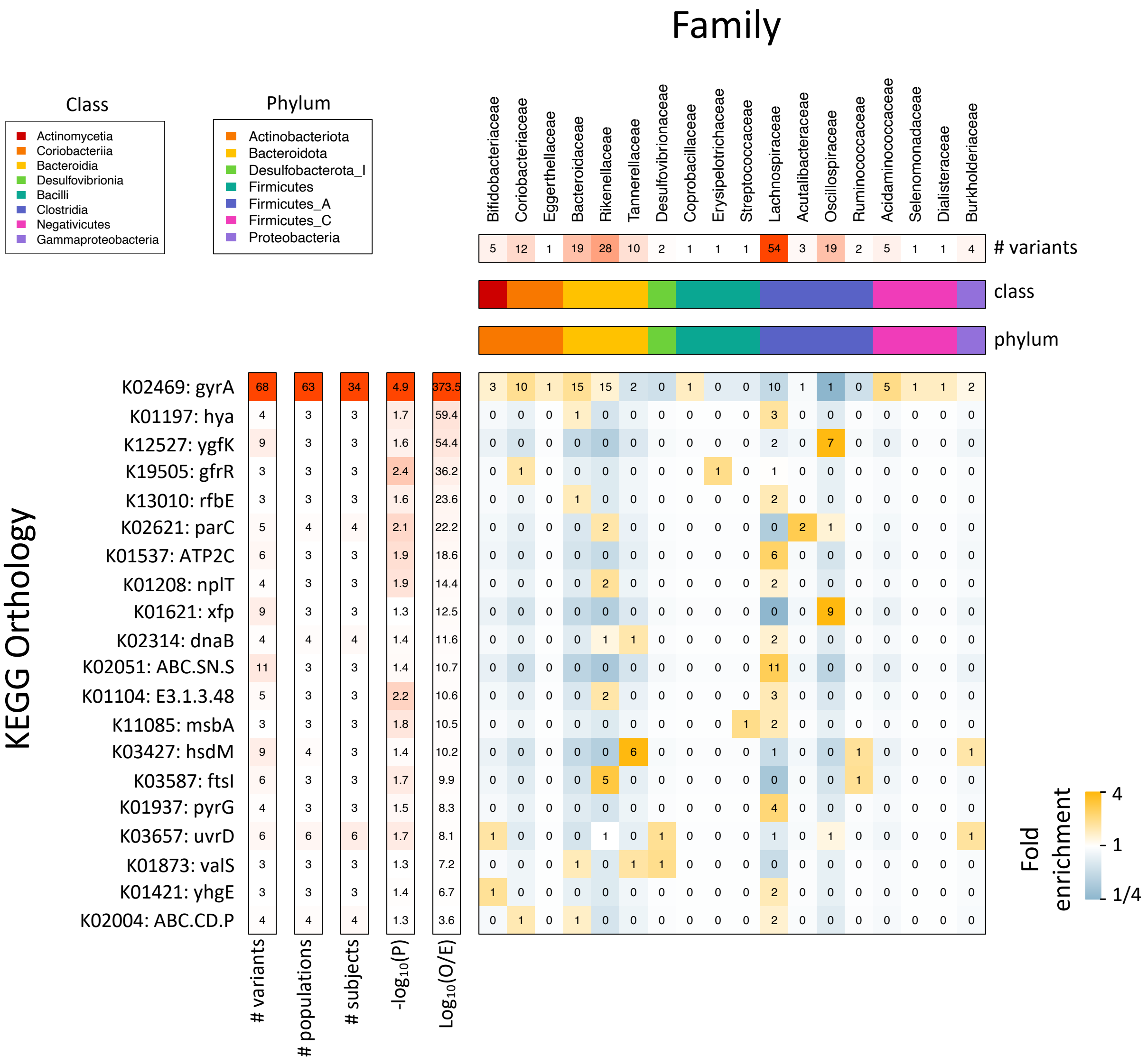


**Fig. S14. KEGG orthology and taxonomic identity.** Variants associated with a significantly enriched KO were stratified by KO (y-axis) and by taxonomic family (x-axis). Shown for each KO (left to right) is the number of associated variants, number of populations, number of subjects, a log-transformed transformed *p*-value, and the enrichment ratio. Shown for each family (top to bottom) is the number of associated variants, the class and the phylum (color legends in top left). The matrix (middle) shows the observed number of variants for each KO-family combination, colored according to fold enrichment of the observed number of variants over the number expected if KOs and families were independent.

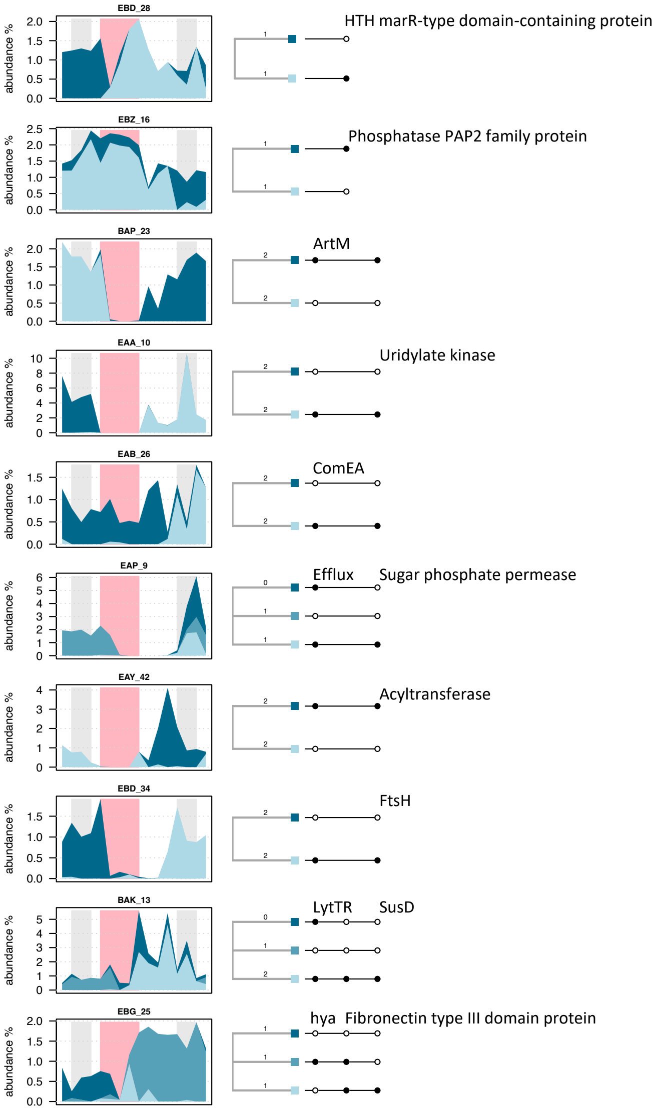


**Fig. S15. Examples of evolving populations not associated with mutations in *gyrA*.** We focused on populations that had a resolved *gyrA* gene without dynamic variants and sweeping variant associated with a non-synonymous substitution in genes other than *gyrA*. Shown are the 10 populations with the smallest number of dynamic variants. See Supp. Figure S13 for a description of the figure components. Associated gene annotations, assessed through KO, Uniref100 and PFAMs, are above each example. All sweeping variants except for a non-synonymous substitution in a hyaluronoglucosaminidase (K01197) in population EBG_25 were idiosyncratic, i.e., their associated KO was not significantly associated with parallel evolution in our study. See Supp. Table S2 for a complete description of variants.

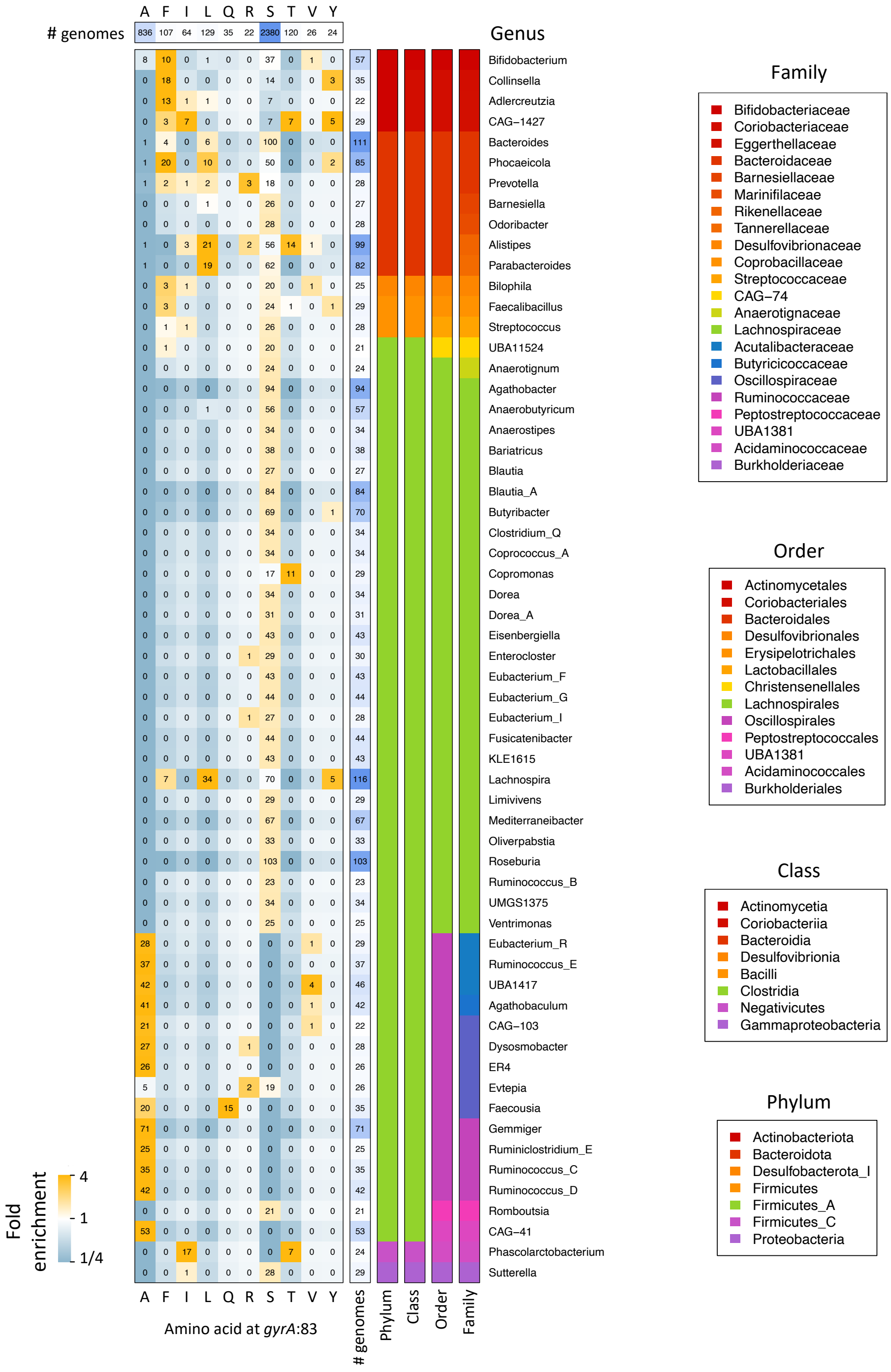


**Fig. S16. *gyrA*:83 baseline amino acids and taxonomic identity.** The amino acid at *gyrA*:83 of *gyrA* was resolved for 3896 genomes. Genomes were stratified by amino acid (top) and genera (right). The matrix (middle) shows the observed number of genomes with *gyrA*:83/genera combination, colored according to fold enrichment of the observed number of genomes over the number expected if taxa and amino acid identity were independent. *Clostridia* families are split between the use of serine (e.g., *Lachnospiraceae*) and adenine (e.g., *Oscillospiraceae*). *Actinobacteria* and *Bacteroidota* combine serine and other amino acids.

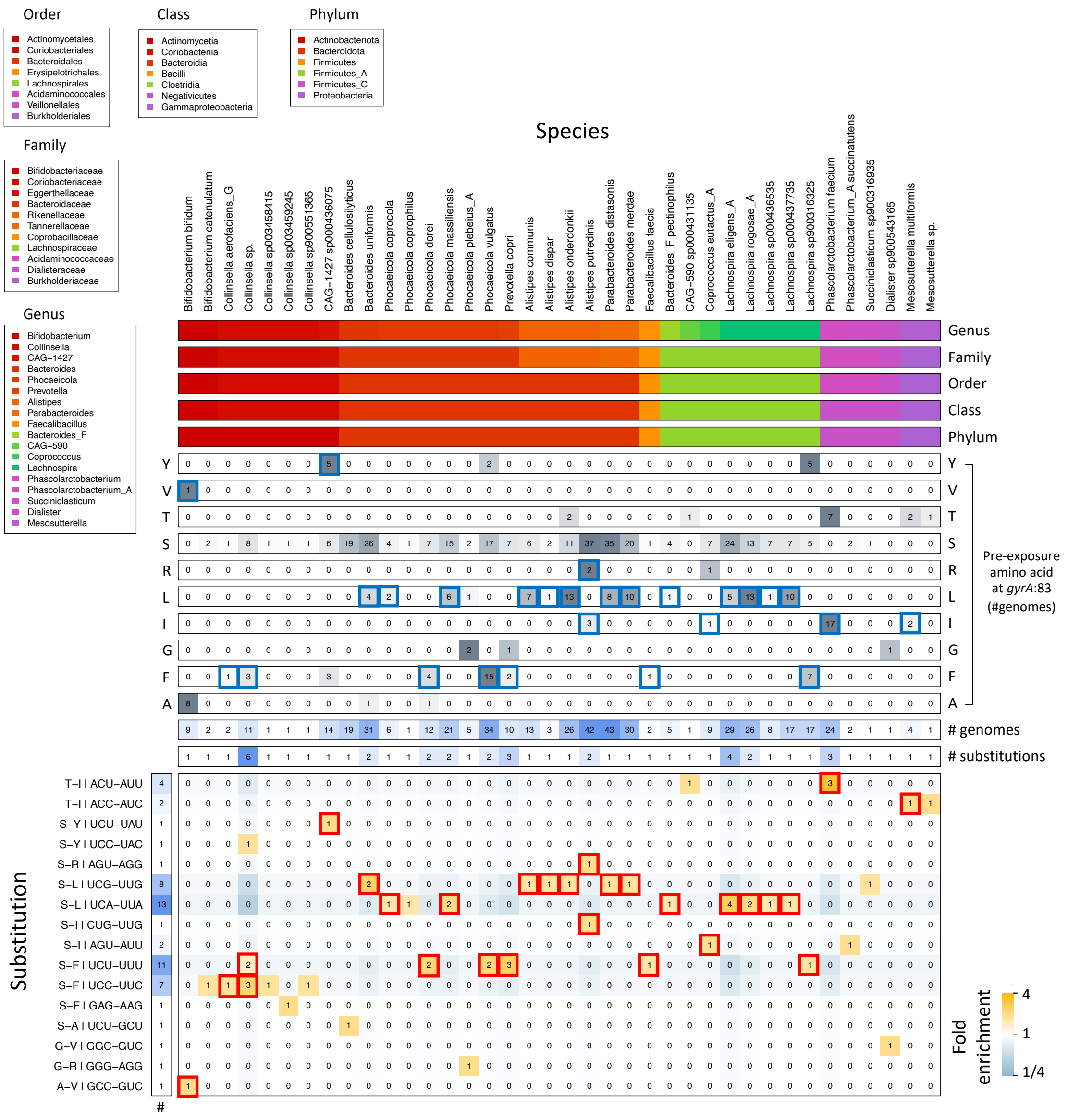

**Fig. S17. Substitutions and taxonomic identity.** Shown are 56 substitutions at *gyrA*:83, stratified by source and target codons (left) and species (top). For each species, shown are the number of genomes with specific pre-exposure amino acids at *gyrA*:83, total number of genomes and number of genomes with a substitution. The matrix (bottom) shows the observed number of substitutions for *gyrA*:83/genera combinations, colored by enrichment (as in Supp. Figure S16). There were 26 species in which the substituting amino acid (highlighted with a red rectangle) was present prior to the exposure (highlighted with blue rectangle). For example, *Bifidobacterium bifidum* had a single substitution to valine during exposure (A83V) and valine was also present before the exposure (1/9 genomes). The association between substituting and pre-existing amino acids indicate baseline amino acid distributions may reflect prior exposure and resistance. *Colinsella sp.* represents unclassified species of the *Colinsella* genus.

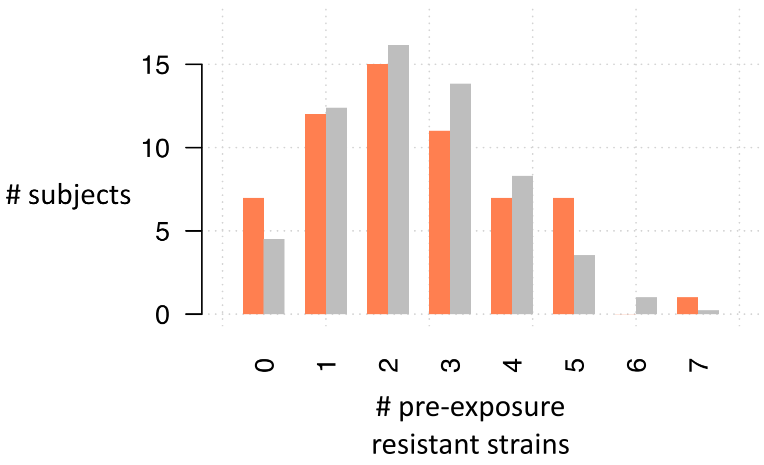


**Fig. S18. Pre-exposure resistant strains across subjects.** The number of resistant populations, defined as populations with a species-specific *gyrA*:83 resistant allele prior to the ciprofloxacin exposure, were counted per subject. Shown in orange is the number of subjects (y-axis), as a function of the number of resistant populations (x-axis). Shown in gray is a background model, generating through n=10,000 permutations of the data that randomly shuffled *gyrA*:83 alleles between subjects while respecting species. The Real and permutated distributions of the number of resistant populations per subject were not significantly different, based on an asymptotic two-sample Kolmogorov-Smirnov test.

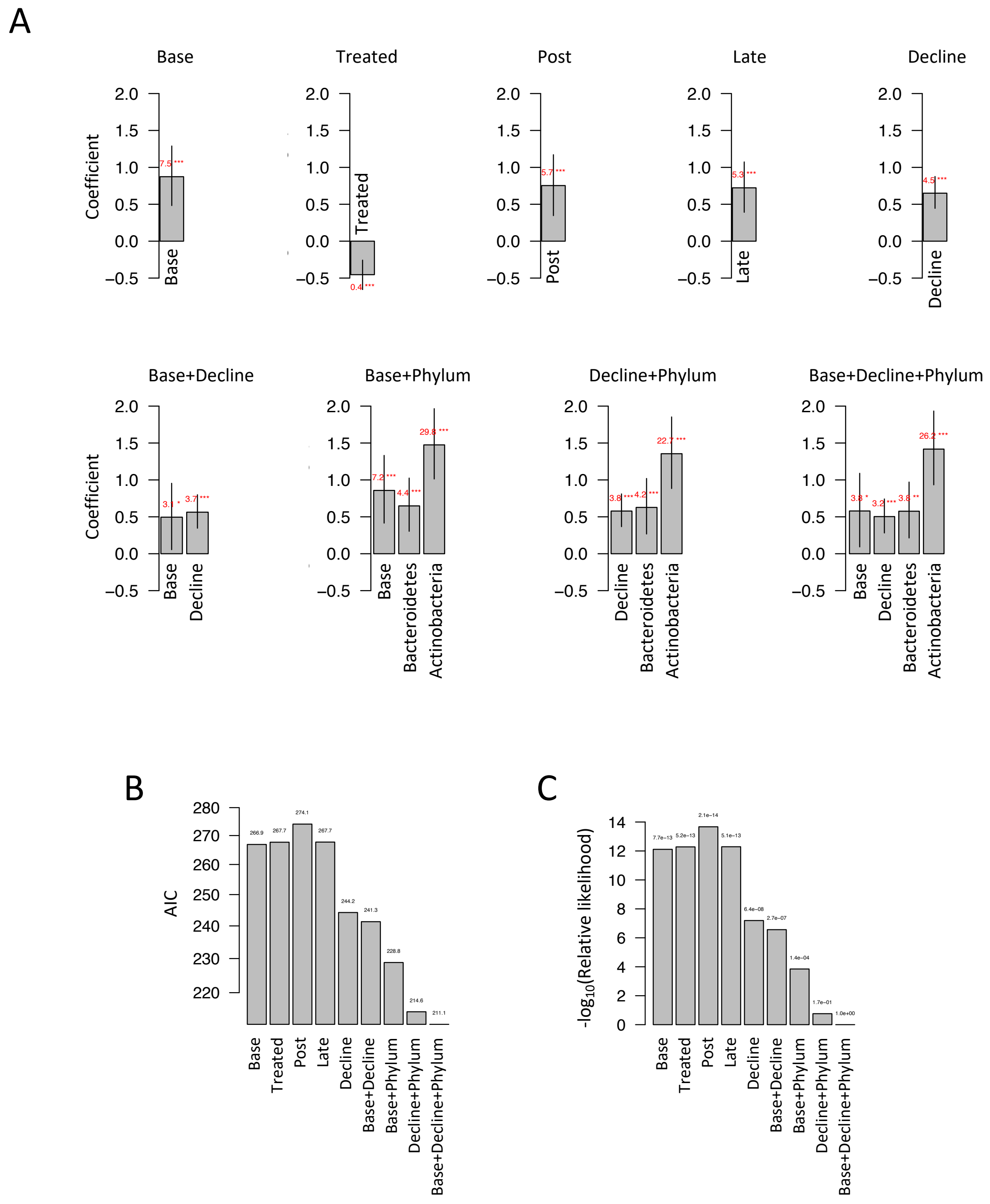


**Fig. S19. *GyrA* evolvability models.** Comparison of 9 logistic regression models with *gyrA* evolvability as the response variable. A) Coefficient values (y-axis, $log_{10}$ scale) for all coefficients (x-axis). Exponentiated coefficient values depicted in red text. *p*-value significance represented with asterisks: P<0.001 (***), P<0.01 (**), P<0.05 (*). Confidence intervals (5% to 95%) depicted with vertical lines. B) Akaike information criterion (AIC, y-axis), for all models (x-axis). Values depicted in text above bars. C) The relative likelihood (y-axis, log-transformed) for all models (x-axis). Values depicted in text above bars.

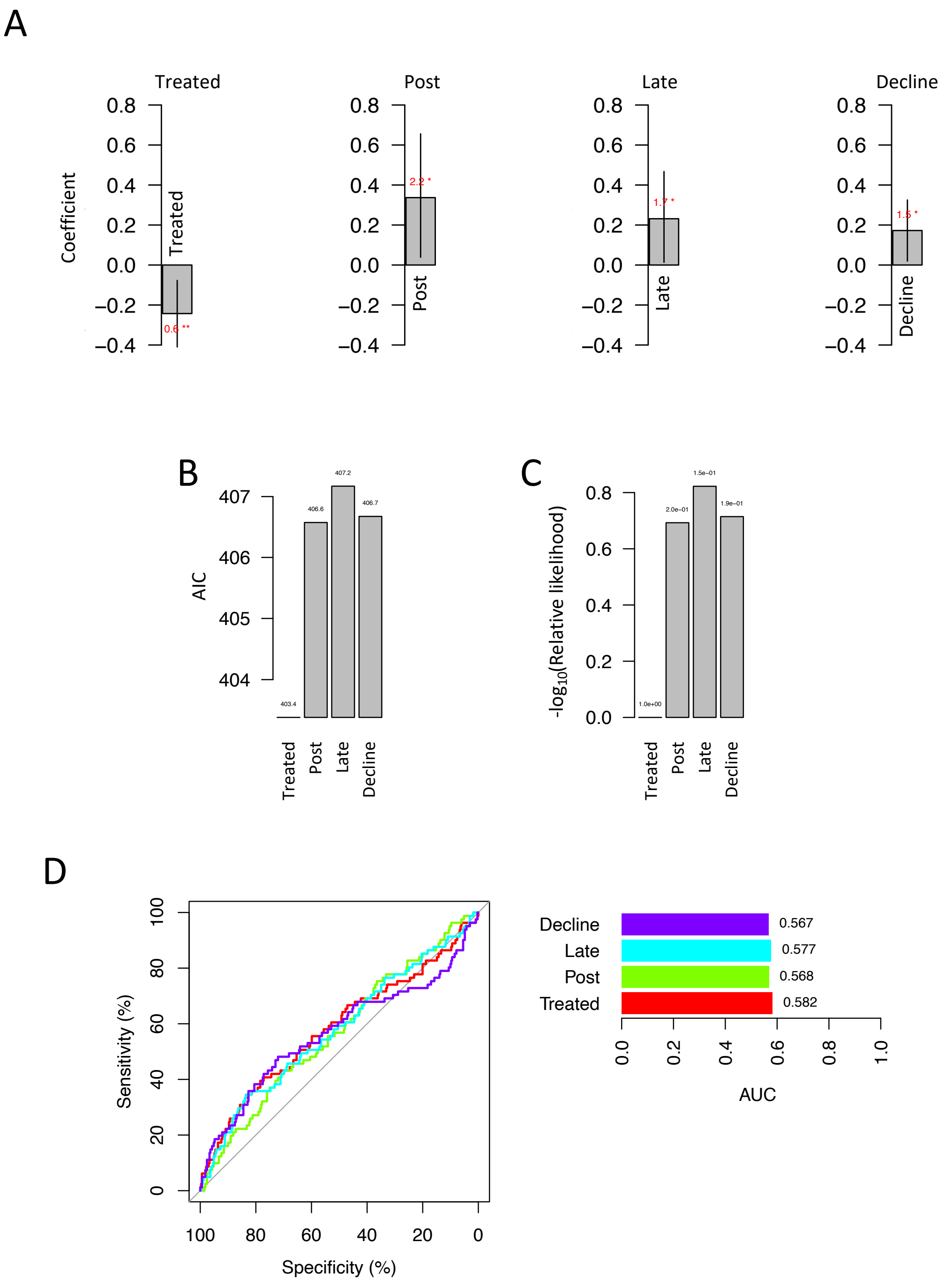


**Fig. S20.** ***Non-GyrA*** **evolvability.** The 9 models described in Supp. Figure S19 were fitted with non-*gyrA* evolvability as the response variable. Shown are 4 models for which the coefficients were significant (*p*-value below 0.05). A) Model coefficients. B) AIC comparison. C) Relative likelihood comparison. D) ROC curves and AUC values of significant models. See Supp. Figure S19 for the legends of panels A-C and Figure 4D for the legend of panel D.

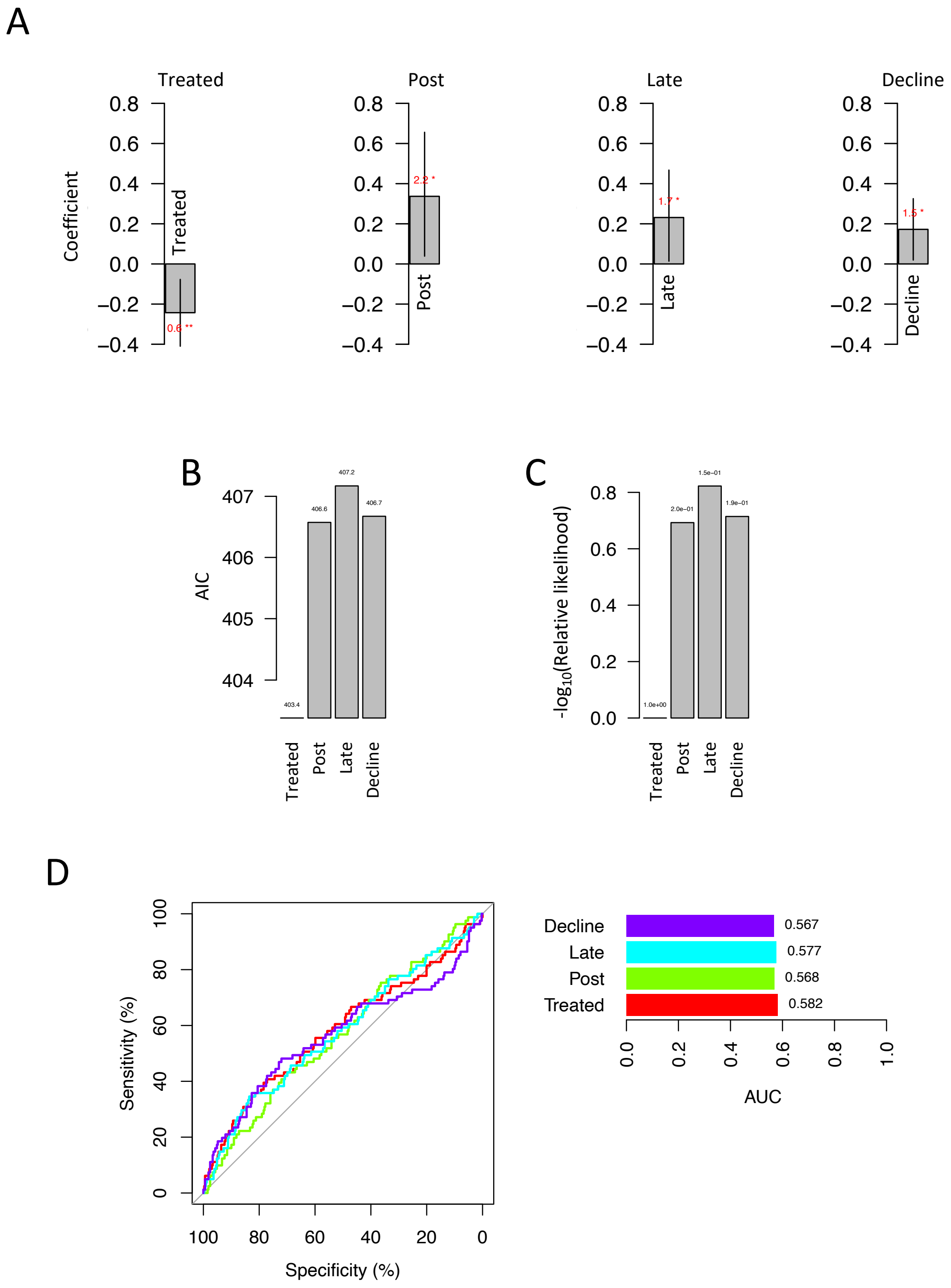


**Fig. S21. Evolving populations across subjects.** The number of evolving populations (involving *gyrA* sweeps on top, any sweep on bottom) were counted per subject. Shown in orange is the number of subjects (y-axis, numbers above bars), as a function of the number of evolving populations (x-axis). Shown in gray is a background model, generating through n=1000 permutations of the data that randomly shuffled populations between subjects while respecting genera. The Real and permutated distributions of the number of evolving populations per subject were not significantly different, based on an asymptotic two-sample Kolmogorov-Smirnov test.
